# Mitotic Chromosome Condensation Driven by a Volume Phase Transition

**DOI:** 10.1101/2021.07.30.454418

**Authors:** Andrew J. Beel, Pierre-Jean Matteï, Roger D. Kornberg

## Abstract

Procedures were devised for the reversible decondensation and recondensation of purified mitotic chromosomes. Computational methods were developed for the quantitative analysis of chromosome morphology in high throughput, enabling the recording of condensation behavior of thousands of individual chromosomes. Established physico-chemical theory for ionic hydrogels was modified for application to chromosomal material and shown to accurately predict the observed condensation behavior. The theory predicts a change of state (a “volume phase transition”) in the course of condensation, and such a transition was shown to occur. These findings, together with classical cytology showing loops of chromatin, lead to the description of mitotic chromosome structure in terms of two simple principles: contraction of length of chromatin fibers by the formation of loops, radiating from a central axis; and condensation of the chromosomal material against the central axis through a volume phase transition.

**One sentence summary:** The mitotic chromosome is an axially scaffolded ionic hydrogel, undergoing a volume phase transition to achieve a condensed state.

---

Eukaryotic chromosomes undergo several gross structural rearrangements during nuclear division: they axialize (adopting a cylindrical form), they contract length-wise, and they condense. Axialization is thought to depend on the formation of a central proteinaceous “scaffold,” which has been convincingly demonstrated for meiotic chromosomes but remains contentious in the case of mitotic chromosomes. Length-wise contraction is thought to be explained by the formation of arrays of chromatin loops, radiating from the central axis. By contrast, the physical basis for condensation has remained obscure, as have its cellular mediators.

The existence of a scaffold in the center of the mitotic chromosome was proposed long ago on the basis of electron micrographs of chromosomes depleted of most proteins, dehydrated in ethanol, and stained with uranyl acetate (*1*). Biochemical analysis of the scaffold suggested a rather simple composition, dominated by two proteins (*2*), later identified as topoisomerase II (*3*) and condensin (*4*). The concept has received recent support from chromosome conformation capture studies of mitotic cells (*5*). Others, however, have argued against the existence of a scaffold (*6, 7*). Indeed, while proteinaceous cores are readily apparent in meiotic chromosomes (*8*), extensive analysis by thin-section electron microscopy has failed to reveal a corresponding core in mitotic chromosomes (*6*). The central structure, visible only by immunofluorescence or upon denaturation, may simply reflect the axial concentration of putative scaffold components.

From the chromosome’s central axis (proteinaceous or otherwise) radiate loops of chromatin, as demonstrated by classical cytologic studies of lampbrush chromosomes (*9*) and subsequently by electron microscopic studies of histone-depleted mitotic chromosomes (*1*). The formation of such loops at least partly explains the reduction in chromosomal length during nuclear division. Loops are thought to arise through a process of “extrusion,” catalyzed in an ATP-dependent fashion by condensin complexes (*5, 10, 11*).

Beyond length-wise contraction, mitotic chromatin is also condensed in volume, achieving a density of 25 percent or greater (*12*), comparable to the density of protein crystals. The basis for this effect is unknown. Although the formation of loops achieves linear contraction, it is an inherently isopycnic process, exchanging length for width and leading to no increase in the density (viz. condensation) of the chromosomal material. Condensins, despite their name, are apparently not involved, because abrupt, auxin-mediated destruction of condensin in mitotic cells does not prevent condensation; in the absence of condensin, the entire chromosomal material condenses into a single compact mass (*13*). *Linear contraction and condensation are evidently distinct processes*.

We sought to understand the basis for chromosome condensation. Our approach consisted in analyzing isolated mitotic chromosomes under various solution conditions with the use of procedures developed to quantify condensation behavior. The results were interpreted in terms of the theory of polyelectrolyte gels, giving insight into the basis for chromosome condensation.

## Controlled chromosome decondensation and recondensation *in vitro*

It has long been known that chromosomes expand and contract upon lowering and raising the ionic strength of the solution (*14, 15*). This behavior has been proposed to reflect the interconversion of chromatin between 10-nm and 30-nm fibers (*16*), as occurs for isolated chromatin subjected to similar ionic shifts (*17*). While the physical chemistry of DNA and chromatin condensation have been studied (*18–20*), a detailed physico-chemical analysis of the condensation of native chromosomes, supported by quantitative observations, is wanting. Limitations of prior work include a reliance on qualitative techniques, the observation of small numbers of specimens, poor demonstration of chromosome morphology, the application of condensing agents over narrow ranges of concentration, and ill-defined perturbations due to the mode of reagent delivery. We sought to overcome these limitations by the quantification of morphologies of individual chromosomes as a function of environmental variables in a high-throughput manner. We immobilized chromosomes within an optically interfaceable, parallel-plate flow cell, and after conditions for the stable adsorption of chromosomes to the objective-proximal surface of the flow cell were established (cf. Supplementary Information), chromosomes were maintained in precisely defined environments by perfusing solutions of various composition.

By means of this experimental system, the condensation state of chromosomes could be precisely adjusted over a wide range (Figure 1A). An advantage of this system is that, in the event of associative interactions between chromosomes and components of the solution, the free concentrations of those components are known, because perfusion is continued until chromosome morphology ceases to change. Reduction of the ionic strength to approximately 5 *×* 10*^−^*^3^ (molar basis) resulted in a ten-fold increase in chromosome volume (Figures 1B and S1A); in this state, water accounted for at least 90% of the volume of a chromosome. Chromosomes could be further expanded by further reduction of the ionic strength (see below).

**Figure 1:**
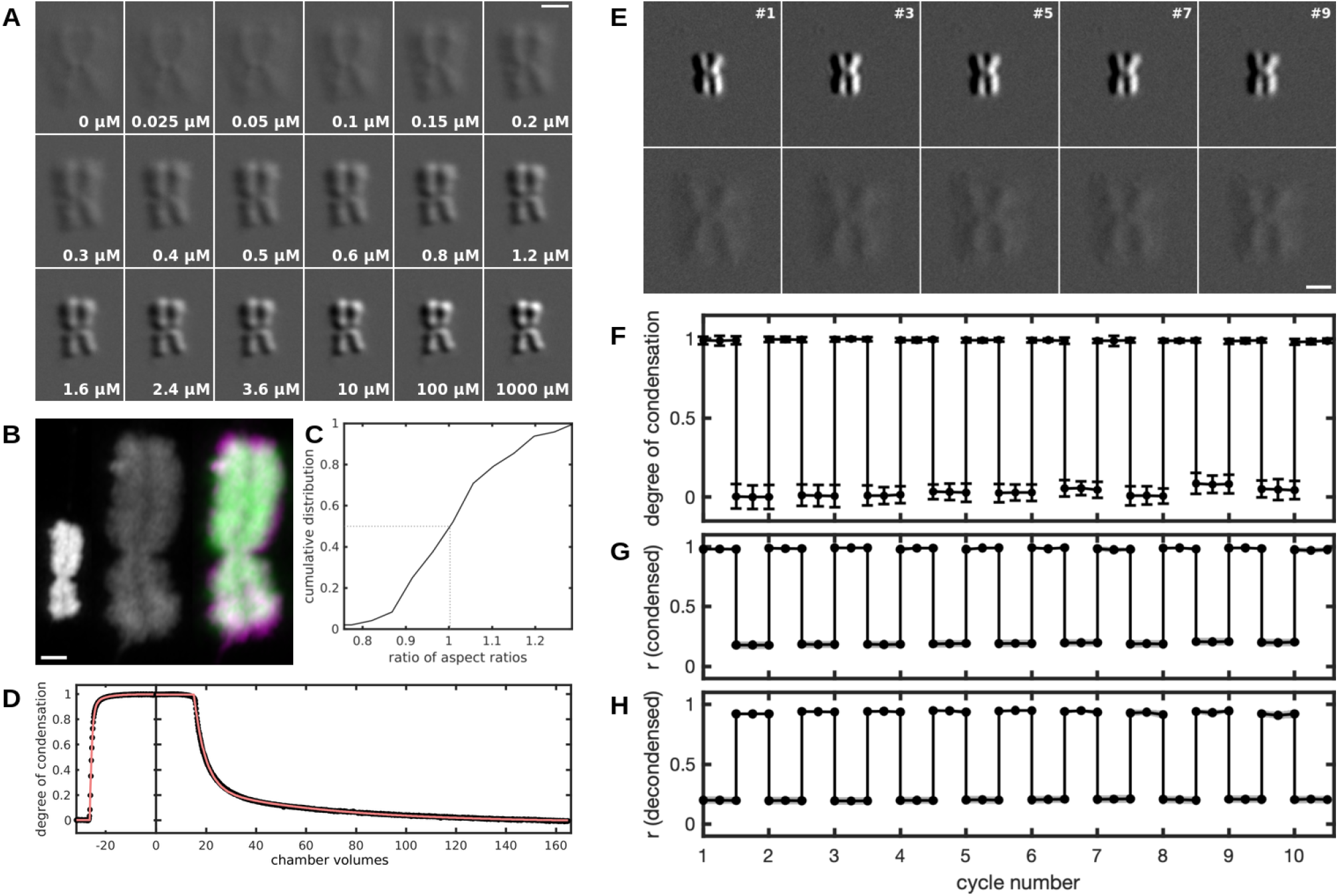
Controlled chromosome decondensation and recondensation *in vitro*. (A) DIC images of a chromosome demonstrating precise tuning of the degree of condensation by varying the composition of the perfusate. Solutions consisted of 5 mM Tris-HCl (pH 7.5), 2 mM KCl, and varying concentrations of spermine (indicated on the panels). Scale bar, 2 µm. (B) Maximum intensity projections of a chromosome stained with Sytox Green (10 nM) in the presence (left) or absence (middle) of 375 µM spermidine, showing variation in chromosome volume over one order of magnitude. The intensity of the middle chromosome was enhanced two-fold to aid visibility. The decondensed chromosome is shown again in the right panel (magenta) in superposition with the condensed chromosome (green) after the latter was isotropically expanded *in silico*. Scale bar, 1 µm. (C) Cumulative distribution of the ratio of aspect ratios measured in three spatial dimensions for condensed and decondensed states of *n* = 16 chromosomes. The midpoint of the distribution occurs at a ratio of unity, indicative of isotropic expansion. (D) A kinetic study of the condensation and decondensation of *n* = 161 chromosomes prepared in a decondensed state and then exposed to 375 µM spermidine at a rate of one chamber volume per minute (left). Decondensation was subsequently effected by flowing a spermidine-free solution [5 mM Tris-HCl (pH 7.5), 2 mM KCl] at the same rate (right). Dots represent data points and the line a biexponential model. (E) DIC images of a chromosome that was alternately condensed and decondensed by sequential provision and removal of 375 µM spermidine. Each column of the montage corresponds to an odd-numbered cycle (first, third, fifth, seventh and ninth), and the rows correspond to 375 µM (top) and 0 µM (bottom) spermidine. Scale bar, 2 µm. (F) Average condensation profile over several condensation-decondensation cycles for *n* = 140 chromosomes; error bars denote ±1*σ*. (G) Pearson correlation coefficient between each image of a chromosome obtained during repeated condensation-decondensation cycling (convolved with a 1*σ* Gaussian kernel) and the average of all images of the condensed states of that chromosome obtained during cycling (averaged over *n* = 140 chromosomes; shading represents the 95% confidence interval of the estimate of the mean). (H) Same as panel G but for correlation with the average image of the decondensed state.

Swelling and deswelling of chromosomes reflect the decondensation and recondensation of chromatin, because mass density observed in phase contrast micrographs corresponded with chromatin density observed by fluorescence microscopy of chromosomes bearing a fusion of histone 2B and green fluorescent protein (H2B-GFP) (Figure S2A). This behavior is not specific to the state of chromatin in mitotic chromosomes, as isolated nuclei exhibited similar behavior (Figure S2B).

Chromosomes were previously swollen in solutions of elevated ionic strength (“reentrant” swelling, discussed below), and the length and width of the chromosomes were shown to vary proportionately, leading to the suggestion that swelling was isotropic (*16*). This result was not conclusive because the analysis was two-dimensional — the third dimension, the thickness of the chromosomes, was not determined. We obtained unambiguous evidence for isotropy by subjecting three-dimensional images of condensed chromosomes to isotropic expansion *in silico* (Figure 1B; green) and superimposing the results upon three-dimensional images of their decondensed states (Figure 1B; magenta). The Pearson correlation coefficient between the calculated and observed three-dimensional images was 0.957 *±* 0.023 (*n* = 54), demonstrating three-dimensional isotropy (Figure S1B). The reciprocal transformation (*in silico* compaction of decondensed chromosomes) gave identical results. Further support for isotropy came from an analysis of aspect ratios computed along cyclically permuted dimensions (Figure S1C), and from the centering about unity of the distribution of the ratio of aspect ratios in condensed and decondensed states (Figure 1C). The demonstration of three-dimensional isotropy was important for our subsequent studies, to assure that the adsorption of chromosomes on the surface of the flow cell did not interfere with decondensation and recondensation behavior.

Because intercalating dyes invariably caused chromosome condensation (Figure 2A), and because photobleaching of fluorescently labeled histone proteins limited the period of observation, chromosomes were imaged as phase objects by differential interference contrast (DIC) microscopy. Computational procedures were devised for the extraction of chromosomal shapes and dimensions from DIC micrographs (Figure 2B), allowing for thermodynamic (Figure 2B, bottom left) and kinetic (Figure 1D) analyses of chromosome condensation. The results of the computational procedures were validated by manual measurements (Figure S3). The maximal and minimal degrees of condensation were operationally defined with respect to solutions promoting nearly complete condensation or decondensation (5 mM Tris and 2 mM KCl at pH 7.5, with or without 0.375 mM spermidine, an essential polyamine known to condense DNA and chromatin (*21, 22*)). The swelling dynamics of thousands of individual chromosomes were quantitatively characterized in this way.

**Figure 2:**
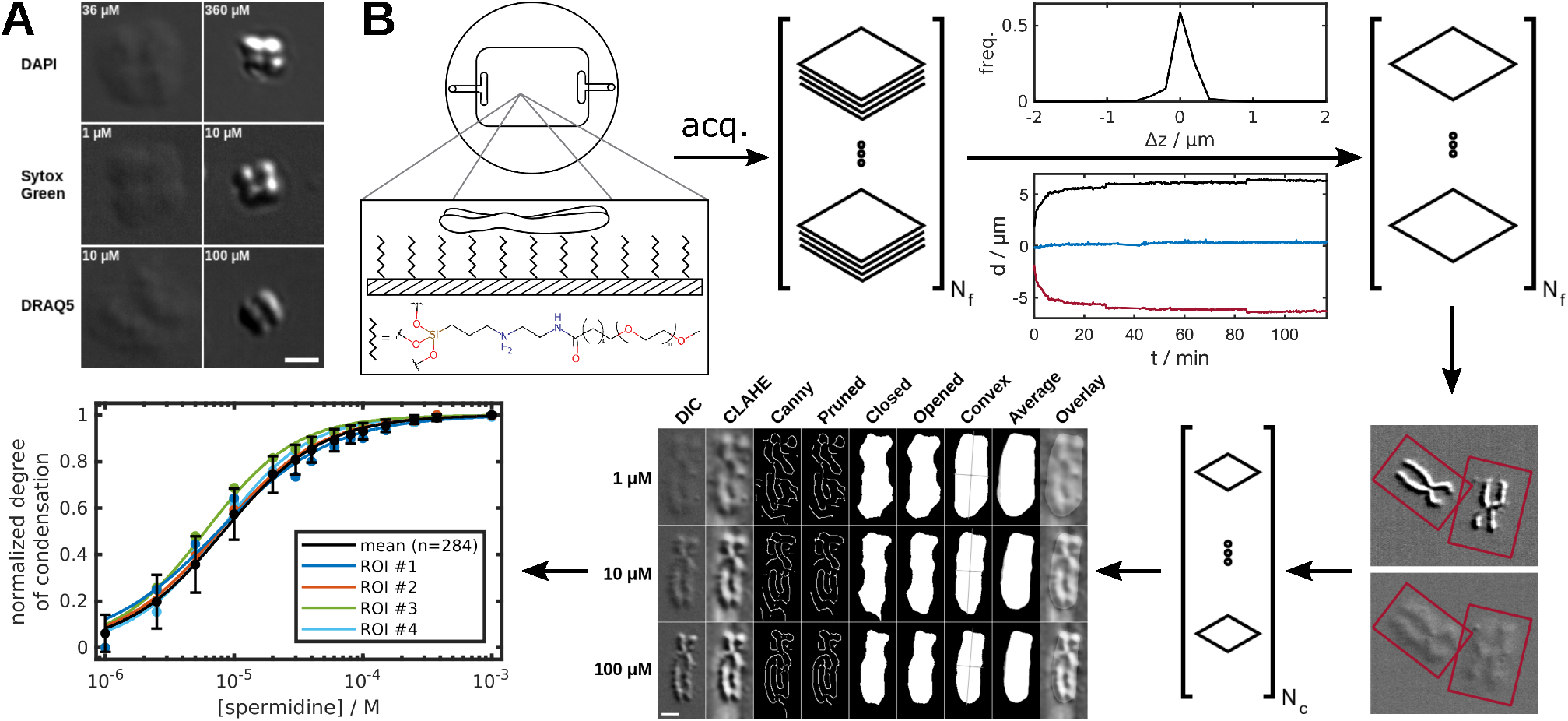
Development of high-throughput methods for minimally perturbative characterization of chromosome morphology. (A) Fluorescent dyes perturb morphology by inducing chromosome condensation. Chromosomes were immersed in the indicated concentrations of dye, mounted on coverslips, and imaged under DIC optics; scale bar, 2 µm. (B) Overview of the experimental approach developed herein. An optically interfaceable, parallel-plate flow cell (top left), supporting immobilized chromosomes, was employed. A chromosome is shown adsorbed to a layer of methoxy-PEG-3-(2-aminoethylamino)propyltriethoxysilane (mPEG-AEPTES). Multiple fields (*N_f_*) of chromosomes are imaged in three dimensions (denoted by stack of planes) in multiple states of condensation (denoted by ellipsis). The images are subjected to automatic, three-dimensional drift correction. The Δz histogram shows good agreement between manually and automatically determined focal planes. Shown below is an example of drift in *x* (blue) and *y* (red), as well as overall in-plane motion (black). Applying these corrections affords a series of aligned and focused slices (depicted in the top right). Single chromosomes are selected (red boxes), giving *N_c_* chromosomal regions of interest (ROIs) (for a typical experiment, *N_c_* is 100–1000). Each ROI is subjected to morphological analysis (bottom middle: rows represent a single chromosome in three states of condensation, obtained by bathing in the indicated concentrations of spermidine; columns correspond to sequential processing steps), yielding condensation profiles (bottom left) representing the average behavior of many individual chromosomes. Morphological image processing steps comprise the following: (DIC) Micrographs (scale bar, 2 µm) subjected to linear contrast adjustment with limits determined by the minimum and maximum pixel intensities of the series of images for a given chromosome. (CLAHE) DIC images convolved with a Gaussian kernel and enhanced by contrast-limited adaptive histogram equalization. (Canny) Edge map resulting from the Canny algorithm. (Pruned) Refined edge map obtained by removal of short edges with an efficiency related to their proximity to the image perimeter. (Closed) Closure of the pruned edge map with a large, disk-shaped structuring element. (Opened) Opening of the closed image with a smaller structuring element than the one used for closing, which helps to prune spurs that would otherwise inflate the convex hull. (Convex) The convex hull of the opened image, energy-minimized against the underlying DIC image by the Chan-Vese active contour algorithm with a modest contraction bias. Superposed on the image in gray are the predicted major and minor axes of the chromosome, joined at its centroid. (Average) The preceding steps were repeated for many combinations of parameters and averaged to obtain a representative map. The grayscale value of pixels in the final map indicates the probability that a pixel overlaps a chromosome. (Overlay) Superposition of the probability-weighted mask and the histogram-equalized DIC image. Condensation profiles (bottom left) show the relationship between the degree of condensation (normalized by the degree of condensation in a pair of calibration solutions) and some independent variable (spermidine concentration in this example). Profiles are shown for individual chromosomes (ROIs 1–4) and as an average of many chromosomes (black curve).

As noted above, observations of chromosome decondensation and recondensation have been made in the past (*14–16*), but quantitative evidence for morphological reversibility over repeated cycles is lacking. We also observed that chromosomes could be repeatedly decondensed and recondensed with qualitative preservation of morphology (Figure 1E), and we applied our computational procedures for quantitative analysis. The degree of condensation was the same for the condensed state at every cycle and for the decondensed state as well (Figure 1F). To quantitatively assess the recovery of chromosome morphology from cycle to cycle, Pearson correlation coefficients were computed between images of chromosomes acquired during each cycle and the average images of those chromosomes in condensed (Figure 1G) and decondensed (Figure 1H) states. The Pearson coefficient among all pairwise comparisons of the images of a chromosome in a condensed state obtained during the course of multiple cycles, averaged over all chromosomes, was 0.983 *±* 0.022; the corresponding value for pairwise comparison of images of decondensed chromosomes was 0.945 *±* 0.033. In contrast, images of condensed and decondensed chromosomes were uncorrelated (*r* = 0.198 *±* 0.118). We conclude that to the resolution of our analysis, *in vitro* chromosome decondensation is reversible (at the end points of the forward and reverse processes — see below), and that the structure following recondensation is reflective of the native state.

## Chromosomes as axially-scaffolded ionic hydrogels

The decondensation of chromosomes in low salt can be attributed to the Donnan effect, whereby the abundance of negative charges of the DNA phosphates — which are only half neutralized by histone proteins (*23*) — attract positive counterions to achieve electroneutrality; the greater concentration of counterions inside than outside the chromosome creates an osmotic pressure difference, drawing water into the chromosome. Cation-driven condensation can be explained either by suppression of the Donnan effect through elevated salt concentration, or by neutralization of the negative charge through binding of the cation to DNA (*18*) (or by a combination of both effects). Charge neutralization is known to be important for the condensation of naked DNA and chromatin fragments (*18–20*), but whether other effects are important for chromosome condensation is unknown.

We pursued the question of cation concentration versus cation binding by determining condensation profiles for the family of amine compounds spermine, spermidine, putrescine, and methylamine (which bear nearly four, three, two, or one positive charges at neutral pH). Notably, even the monovalent cation (methylamine) induced chromosome condensation (green points in Figure 3A), indicating a qualitative difference in behavior compared to naked DNA and chromatin. Quantitative analysis of hundreds of individual chromosomes perfused with solutions of these compounds disclosed the nature of the dependence of chromosome condensation on counterion concentration and counterion charge (points in Figure 3A). The data could be fit by binding isotherms (curves in Figure 3A). The logarithm of the effective concentrations exhibited a linear dependence on counterion charge (Figure 3B), and the reactions were non-cooperative (Hill coefficients of *∼*1 for all counterions).

**Figure 3:**
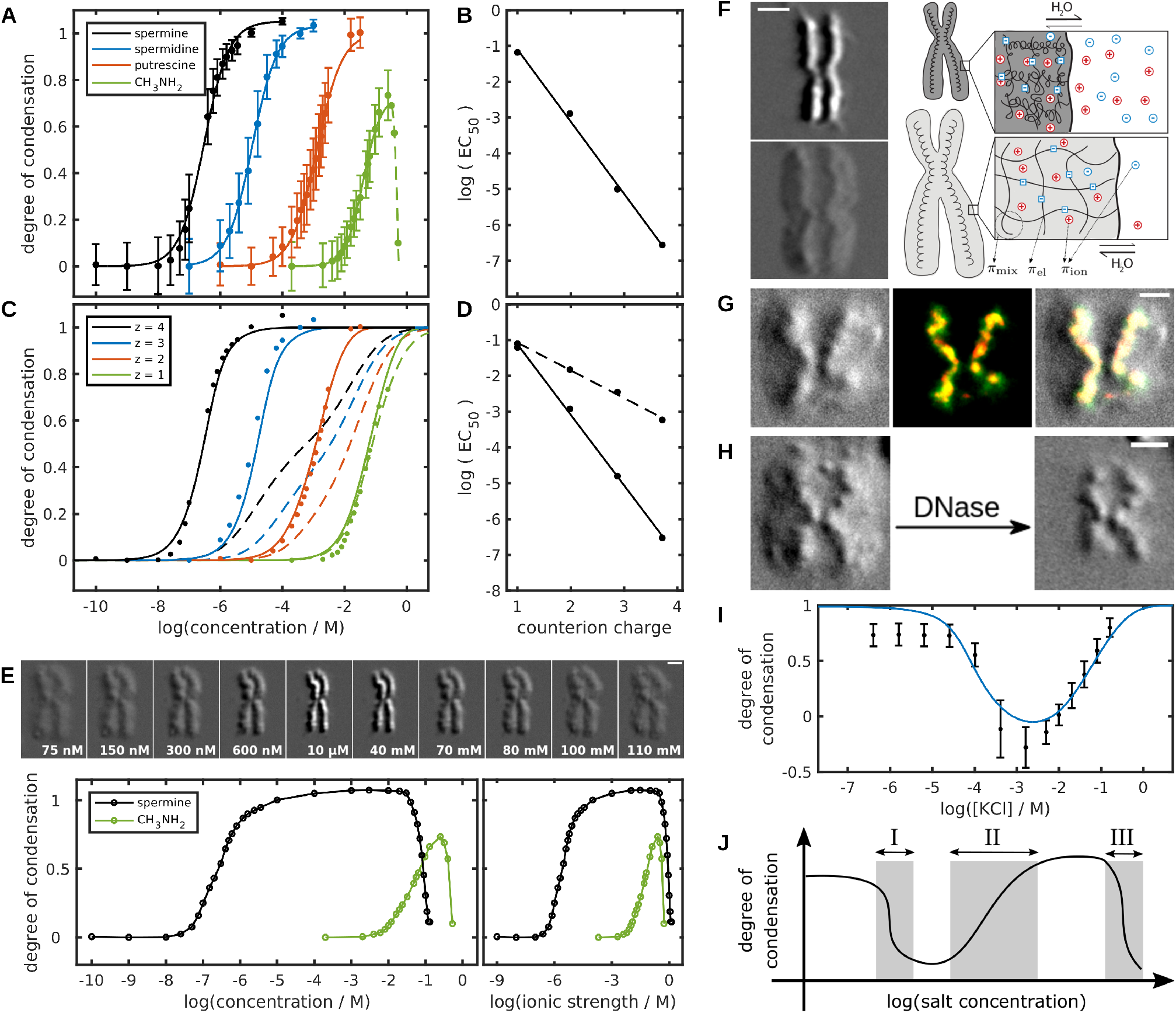
Chromosomes are axially scaffolded ionic hydrogels. (A) Condensation profiles for *n* = 504 (spermine; black), *n* = 367 (spermidine; blue), *n* = 270 (putrescine; orange), and *n* = 208 (methylamine; green) chromosomes are shown with error bars denoting 1*σ*; lines represent leastsquares fits to the Hill model. (B) The concentrations at which condensation is 50% complete (EC_50_) from panel A are plotted with respect to the charge of each ion at pH 7.5, and fit by a straight line (*r*^2^ = 0.996). (C) Theoretical swelling curves for a polyelectrolyte gel without (dashed lines) or with (solid lines) a gel-counterion binding interaction (cf. Supplementary Information). Points represent experimental data from panel A. (D) The EC_50_ of the curves shown in panel C. The solid and dashed lines correspond, respectively, to the sets of solid and dashed curves in panel C. (E) *Top*: Variation of spermine concentration demonstrates reentrancy of chromosome condensation. DIC micrographs are shown for select concentrations; scale bar, 2 µm. *Bottom*: Degree of chromosome condensation for *n* = 205 chromosomes (black), in superposition with that for methylamine (green; copied from panel A), is plotted with respect to the decimal logarithm of concentration (left) or ionic strength (right; calculated in molar units according to 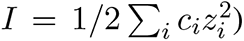. (F) *Left*: DIC images of a chromosome in 375 µM (top) and 0 µM (bottom) spermidine. A central filament is observed upon decondensation. Scale bar, 2 µm. *Right*: Chromosome volume is determined by three pressures, those of mixing (*π*_mix_), network elasticity (*π*_el_), and ion partitioning (*π*_ion_), as illustrated for a chromosome in two states of condensation; at equilibrium these pressures balance. Mixing comprises entropic and enthalpic contributions. Elasticity, empirically supported by chromosome-stretching experiments (*71*), describes the principally entropic resistance to deformation. The entropy of counterion confinement quantifies the electrostatic effects of the polyelectrolyte. Complete equations along with a description of the various quantities can be found in the supplementary material. (G) DIC (left) and fluorescence (middle) images of a decondensed chromosome treated with antibodies recognizing a condensin I subunit, CAP-G (green), and topoisomerase II*α* (red). Areas of overlap are shaded yellow. The merged image (right) demonstrates colocalization of the putative scaffold components with the central filament. (H) DIC images of a chromosome in a decondensed state before (left) and after (right) treatment with DNase I. (I) Chromosomes were titrated with KCl solutions prepared in argon-saturated water. Data (black; represented as mean *σ*) show the degree of chromosome condensation (normalized to [0,1] with respect to calibration solutions). The curve (blue) represents the theoretical predictions from a model parameterized as in panel C, except for the change of background solution from 5 mM Tris, 2 mM KCl to water. (J) The effect of counterions on chromosomes can be divided into three regimes dominated by: (*I*) proton-counterion exchange; (*II*) charge neutralization; (*III*) charge screening.

A fit to binding isotherms does not necessarily mean that condensation is driven by cation binding. All components of the system — chromatin, polyamines, buffer, salt, and water — must be taken into consideration in the interpretation of condensation behavior. We tested the application of a well established theory describing the swelling of a polyelectrolyte gel (*24–26*) which we adapted for chromatin and the components of our system. We calculated the condensation behavior predicted by the theory with parameterization determined by a Monte Carlo minimization procedure (cf. Supplementary Information), from which a subset of the data (that for spermidine-driven condensation) was omitted for the purpose of cross-validation. The results recapitulated the experimental data (points in Figures 3C) when a term for the binding of polyamines was included in the calculations (solid lines in Figures 3C and 3D), but not when this term was omitted (dashed lines in Figures 3C and 3D). Cross-validation analysis demonstrated the predictive strength of the model for data excluded from fitting; specifically, the value of chisquared, reduced by the number of observations, a measure of relative model fidelity, was 0.52 for all data included in the calculations and 0.98 for the omitted subset (spermidine). The theory was predictive of chromosome condensation behavior, *formally identifying the chromosomal material as an ionic hydrogel*.

The three polyamines spermine, spermidine, and putrescine were effective at concentrations orders of magnitude below that required for suppression of the Donnan effect (a concentration comparable to that of unneutralized DNA phosphate, estimated to be on the order of 0.2 M), and their effect was almost entirely explained by binding. The mode of binding, which is not specified by our model, is presumably that of counterion condensation, as shown for the condensation of DNA (*27*) and chromatin fragments (*20*). Contrary to the behavior of multivalent cations, deswelling driven by monovalent cations, exemplified by methylamine (green points in Figure 3C), occurs at a concentration comparable to that of unneutralized DNA phosphate. Furthermore, the predicted behavior was largely unchanged by omission of the term for counterion binding (compare solid and dotted green lines in Figure 3C). Condensation driven by monovalent cations thus reflects suppression of the Donnan effect, with a lesser contribution from binding. Chromosomes adopt an intermediate degree of condensation at physiologic concentrations of monovalent salt (Figure 3A), so more or less condensed states may occur, depending on the conditions of the cellular milieu.

The re-swelling of chromosomes observed with progressively higher concentrations of methylamine (dashed line in Figure 3A), which reproduces the effect of high concentrations of magnesium chloride (*15*), can be interpreted as reentrant swelling, a common attribute of gels (*28*). Spermine induced re-swelling at a lower concentration than methylamine but at the same ionic strength (Figure 3E), demonstrating that, unlike condensation (Figure 3A), which is a function of counterion concentration, reentrant decondensation is a function of the solution ionic strength. Two regimes of counterion concentration were thus apparent: low concentrations, where charge neutralization drives condensation, and high concentrations, where charge screening drives reentrant decondensation. (There is in fact a third regime, whose properties are described below.)

Our calculations (Figure 3C) assume only that chromosomes are polyelectrolyte gels (all lines in Figure 3C) and that chromosomal charge can be neutralized by multivalent cations (solid lines in Figure 3C). The amount of charge borne by a cation, however, is not the only important factor, as quantitative differences were observed for cations having the same amount of charge but different spatial distributions thereof (Figure S5A). Such ion-specific effects are consistent with predictions from atomistic molecular dynamics calculations of the condensation of counterions around a polyelectrolyte (*29*), as well as with differences in the effectiveness of spermidine and hexamminecobalt(III) chloride on DNA condensation (*27*). More highly concentrated charge was associated with greater effectiveness, which could reflect the correlation between smaller ionic radii and shorter DNA persistence lengths (*30*).

Besides neutralization, the effects of multivalent cations on DNA and chromatin have been attributed to other processes such as charge inversion and counterion-induced inter-DNA attraction. These phenomena occur at much higher counterion concentrations than those tested by our experiments; for instance, charge inversion of DNA is induced by *∼*1 mM spermine at an ionic strength of *∼*0.01 (*31*), and attractive inter-DNA forces appear at magnesium concentrations between 10 and 50 mM (*32*). Therefore such effects cannot explain the overall behavior shown in Figure 3A.

The identification of chromosomal material as an ionic hydrogel does not explain the cylindrical shape of chromosomes. A gel formed by the coalescence of polymer chains in the absence of constraints would be expected to be approximately spherical. The breaking of spherical symmetry may be attributed to a central filament, which became apparent upon decondensation of chromosomes (Figure 3F, left). This filament presumably represents the “scaffold” — first demonstrated in electron micrographs as a remnant of denatured chromosomes (*1*), and seen here for the first time in intact form in the native state, without the use of stains. Identification of the filament as the scaffold was supported by immunofluorescence microscopy, which revealed extensive colocalization with putative scaffold components, including condensin I and topoisomerase II*α* (Figures 3G and S4A).

The scaffold could be released from chromosomes by controlled nuclease digestion (Figures 3H and S4B). The scaffold expanded and contracted along with the surrounding chromatin before digestion (Figure S4C), but it lost this capacity after nuclease digestion (Figure S4B), apparently reflecting the transmission of tensile forces from the chromatin. This possibility is further supported by the isotropic nature of chromosome expansion (Figures 1B and S1), implying the application of equal and opposite forces on the scaffold by the expanding chromatin network. Upon removal of chromatin, the scaffold exhibited striking flexibility, undergoing rapid, thermally driven, conformational fluctuations (Movie S1). Such flexibility may be important, for example, for bending of chromosomes at the centromere while under tension from kinetochore-attached microtubule fibers.

On the basis of these observations, the morphology of the mitotic chromosome *may be simply explained by the collapse of a chromatin gel against a central axis*; the electrostatic potential energy of the excess charge, which concentrates at the surface of a gel (*33*), is minimized by cylindrical geometry. The range of condensation states observed over a broad range of cation concentration, irrespective of cation charge (Figures 1A and 3A), need not be ascribed to the interconversion of 10-nm and 30-nm chromatin fibers, as previously assumed (*16*), and can instead be explained simply by a more or less extended configuration of the chromatin fibers (Figure 3F, right).

Partial decondensation of chromosomes also led to the appearance of pairs of centrally located dots, attributed to kinetochores (Figure S4D). Their identity was confirmed by immunostaining with antibodies directed against CENP-A (Figure S4E), a component of the kinetochore inner plate (*34*). Ordinarily invisible by phase imaging, kinetochores were revealed by lowering the refractive index of the chromatin background. Unlike chromatin and scaffold, kinetochores showed no tendency to expand or contract with change of environment (Figure S4F), consistent with their capacity to withstand large forces from the spindle apparatus during chromosome segregation (*35*).

## Confirming the identification of chromosomes as ionic hydrogels

In the theory of polyelectrolyte gels, the principal determinants of gel volume (shown schematically in Figure 3F) are the free energies of mixing, chain elasticity, and ion partitioning (*24*). Derived from these free energies are osmotic pressures which drive uptake or expulsion of solvent; when these pressures balance, the volume of a gel ceases to change. Knowledge of these determinants leads to testable predictions by which the identification of chromosomes as ionic hydrogels can be confirmed. Such predictions include the dependence of swelling behavior on the pH, the ionic strength, the Flory-Huggins parameter, and the relative permittivity of the solvent.

Chromosomes are acidic, and the swelling of an acidic hydrogel is expected to be enhanced in alkaline environments and reduced in acidic ones, an expectation that was confirmed by variation of pH (Figure S5B). The dependence on pH reflects a change in the ionization state of chromatin, but the ionizable groups have not been identified. Inasmuch as the pKa of DNA phosphates is around 0, the pH dependence most likely reflects the ionization state of the histones (*36*) or that of ring nitrogens of adenine and cytosine (*37*). The chromosome-condensing activities of multivalent cations, such as polyamines, and of acids, act through a common mechanism, namely the reduction of chromosomal “fixed charge” density. Reduction is accomplished noncovalently by multivalent cations (through counterion condensation) and covalently by acids (through protonation of chromatin).

A counterintuitive prediction of theory (*25*) — borne out by our calculations for chromosomes (curve in Figure 3I) — is that an ionic hydrogel, which swells upon lowering the ionic strength, will deswell when the ionic strength is lowered further still. Indeed, chromosomes attained a maximally decondensed state at a molar ionic strength of *∼*10*^−^*^3^, from which point *condensation could be achieved by further lowering of the ionic strength as well as by raising it* (points in Figure 3I). Condensation at very low ionic strength derives from an ion-exchange process: as the salt concentration is reduced, hydronium ions are increasingly recruited as counterions. Through protonation of acidic groups, the incoming hydronium ions reduce the fixed charge density of the gel (at an ionic strength determined by the pKa of the gel), thereby depriving the gel of the counterions required for it to swell. Counterion-mediated chromosome condensation can therefore be divided into three concentration regimes, each dominated by a different mechanism (Figure 3J). In the first regime, a proton-counterion exchange process shifts the pH of the chromosomal interior toward neutrality, thereby endowing the chromosome with fixed charge as a result of acid-dissociation reactions. In the second regime, the degree of fixed charge is decreased by an associative process between counterions and charged moieties along the chromosome (with a contribution from the suppression of the Donnan effect which is negligible for all but monovalent cations). In the third regime, there is strong screening of the fixed charge (a short Debye length) and a disruption of chromosome-counterion interactions.

Gel theory also predicts a dependence of the degree of swelling on the thermodynamic quality of the solvent. The impact of solvent quality on the free energy of polymer-solvent mixing is quantified by the Flory-Huggins parameter (*χ*), whose value is determined by the relative energetic strength of heterotypic (polymer-solvent) and homotypic (polymer-polymer and solvent-solvent) interactions. Water, with a sufficient concentration of dissolved ions (Figure 3I), is a good solvent for chromatin. Higher values of *χ*, which favor less swollen states, are expected for less polar solvents. This prediction was confirmed by immersing chromosomes in the polyols glycerol and 2-methyl-2,4-pentanediol: the degree of condensation increased inversely with solvent polarity (Figures S5C and S5D).

The Bjerrum length — the length scale at which the electrostatic energy between two elementary charges reaches parity with thermal energy — is inversely proportional to the dielectric constant. Reduction of the dielectric constant therefore facilitates ion pairing, and is expected to enhance polyamine-driven condensation of chromosomes. To test this prediction, the dielectric constant was varied by mixing water (*ε* = 78.3 at 298.15 K) and glycerol (*ε* = 42.5) in varying proportions. Accordingly, the potency of spermidine increased inversely with the dielectric constant, and was greater by an order of magnitude in nearly pure glycerol compared to aqueous solution (Figure S5E).

## Chromosome condensation as a volume phase transition

Gels are a phase of matter formed by the coalescence of sols and stabilized by some means of cross-linking. In addition to the sol-gel phase transition, gels themselves undergo thermodynamic phase transitions, predicted by classical theory (*38*) and verified experimentally (*39*). Known as volume phase transitions (VPTs), they are observed under certain conditions of the gel and solution.

Hysteresis is a common signature of first-order phase transitions (*40*), and reflects the coexistence of two phases in different proportions along the forward and reverse directions of a phase transition. Hysteresis of gel volume, reflective of a VPT, is predicted by the theory under certain solution conditions: the calculated condensation curves develop an S-shaped inflection, which may resolve as a hysteresis loop because of the thermodynamic instability of states lying along the loop. Consistent with theoretical expectation, hysteresis was observed in condensation-decondensation curves of chromosomes, produced by variation of spermine concentration (Figure S6A) and pH (Figure 4A). The time over which spermine concentration and pH were varied was long in relation to the relaxation time of the system (Figure S6B); the hysteresis loops therefore reflect the occurrence of metastable states (*41*). Chromosome condensation evidently has the properties of a bistable system. The occurrence of a VPT does not require a change in the ionic milieu. At physiologic ionic strength, our calculations predict a VPT driven by an increase in the Flory-Huggins parameter (Figure S7).

**Figure 4:**
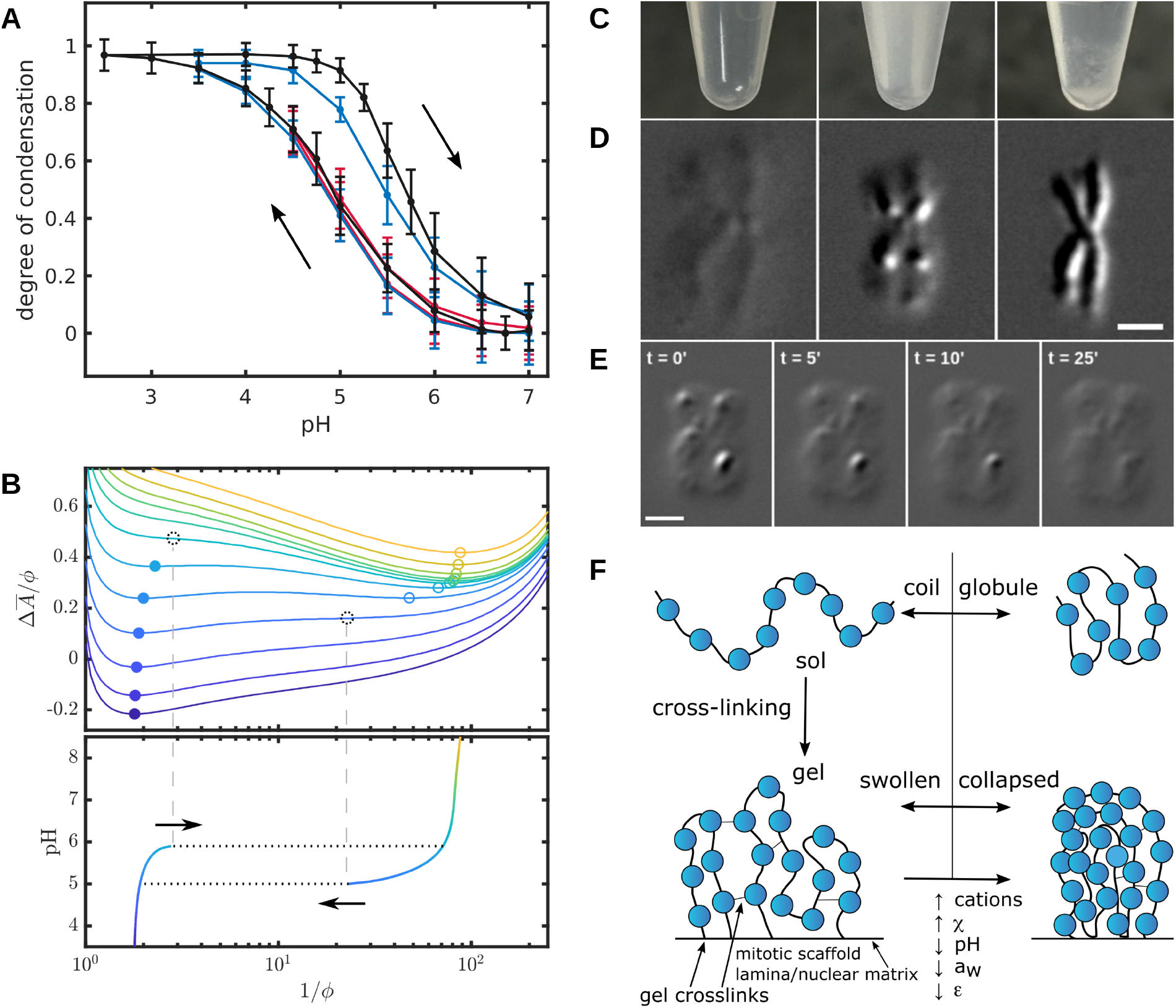
Chromosome condensation involves a volume phase transition of the chromatin gel. (A) Hysteresis caused by variation of pH. Chromosomes were condensed by acidification and subsequently decondensed by alkalinization of the solution (path indicated by arrows). Hysteresis was not observed for low limiting degrees of condensation (red; *n* = 580), but emerged as the limiting condensation was increased (blue [*n* = 452] and black [*n* = 147]). Error bars denote ±1*σ*. (B) Re<u>la</u>tionship between phase transitions, hysteresis, and bistability. *Top*: The Helmholtz free energy (Δ*A*) divided by the polymer volume fraction (*φ*) is plotted with respect to the reciprocal of the polymer volume fraction. (Higher values of *φ* correspond to more condensed states.) Free energy landscapes are shown for various solution pH, indicated by color (purple: pH 3.5; orange: pH 9.5). The sharp rise of free energy at the left and right sides of the graph is attributable to the entropies of demixing and chain extension, respectively. Local minima are indicated for condensed states (closed circles) and decondensed states (open circles). A single minimum is found at more extreme pH, whereas two are found at intermediate pH. Dotted black circles represent extrapolations of the trajectory of the local minima into unstable territory; such states would immediately relax to the opposite state (i.e., from condensed to decondensed or vice versa). Metastability is due to the persistent occupation of a local minimum under conditions where it is no longer the global minimum (e.g., the circles preceding the dotted ones). As the local maximum becomes less prominent, the probability that thermal fluctuations lead to escape increases, and the gel may undergo a phase transition to the state corresponding to the global minimum. The pH at which such a transition occurs differs for the forward and reverse reactions because of the asymmetry of the underlying free energy functions. *Bottom*: The solution pH (colored as in the top panel) is plotted with respect to the positions of free energy minima, demonstrating a hysteresis loop. Loop directionality is indicated by arrows. Breaks in the curve (denoted by dotted lines) occur at stationary points (where the first and second derivatives of the free energy are simultaneously zero). Beyond the *closure points* of the loop, there is a single minimum in the free energy (i.e., a horizontal line intercepts the graph only once). As in panel A, a phase transition does not occur for small perturbations, but once one has occurred, return along the same path becomes impossible. (C) Chromosomes condensation was apparent macroscopically by flocculation of suspended chromosomes. A suspension of 0.8 A_260_ units of chromosomes (left) exposed to 375 µM spermidine opacified after one minute at room temperature (middle) and subsequently sedimented under the influence of gravity (right). (D) A chromosome is shown in decondensed (left), condensed (right), and phase-separated (middle) states. (E) Beginning from a phase-separated state (left), chromosomes were washed with 5 mM Tris-HCl (pH 7.5), 2 mM KCl, causing movement of the phase boundary between condensed and decondensed phases until the latter phase was completely eliminated (right). Scale bar for panels D and E: 2 µm. (F) Relation between the condensation of chromatin and chromosomes. Chromatin fibers, present as a dilute solution in water, adopt a coil configuration and behave as a sol (top left); addition of multivalent ions results in polymer collapse through a coil-globule transition (top right) which may be realized intramolecularly or intermolecularly depending on the concentration. These condensates may have the properties of a physical hydrogel (e.g., due to bridging by multivalent counterions), but they will revert to a sol in the limit of infinite dilution. In contrast, cross-linking the polymer (bottom left) through inter-fiber contacts (which may be mediated by cations or proteins) and fiber-matrix contacts (where “matrix” encompasses the scaffold in mitosis and nuclear lamina/matrix in interphase) introduces elastic restoring forces, which prevent dissolution at infinite dilution. Under certain conditions, the gel will collapse to a condensed state (bottom right). This VPT of the chromosomal gel (bottom) is a manifestation of the coil-globule transition of the chromatin sol (top). This transition is responsive to environmental parameters, as indicated.

Hysteresis of the chromosomal material can be understood through an analysis of the free energy of a chromosome modeled according to polyelectrolyte gel theory. The dependence of the free energy on the polymer volume fraction shows local minima for condensed and decondensed states (filled and open circles, respectively, in Figure 4B, upper panel). These local minima shift as the pH is raised or lowered (trajectories of filled and open circles), and ultimately disappear at different points for condensation and decondensation. Continuation of the trajectories of the minima beyond such points results in unstable states (dotted circles in Figure 4B, upper panel), which will be driven to the opposing state through a phase transition (dotted lines in Figure 4B, lower panel). Because the forward and reverse transitions occur at different pH, a hysteresis loop is formed. Preceding the discontinuities (dotted lines in Figure 4B, lower panel) is a metastable regime, characterized by the presence of two local minima, from which earlier states may be reached by reversal of the system’s trajectory. Alternatively, thermal fluctuations may culminate in a phase transition, after which return along the same path becomes impossible (for more detail, see the legend for Figure 4B). Consistent with this account, much of the swelling process exhibited approximate reversibility of path (red curve in Figure 4A), with the reverse path falling within the scatter of the forward path; beyond a certain point, however, path reversal was no longer observed, resulting in a hysteresis loop (blue and black curves in Figure 4A). Notably, the threshold of irreversibility occurs at a degree of condensation similar to that of chromosomes in physiologic monovalent salt solutions (compare Figure 4A with Figures 3A [green curve] and 3I), indicating that chromosomes may naturally reside in a condensation state near that of the VPT, such that small fluctuations in the ionic milieu could drive a phase transition.

The observation of hysteresis shows that contrary to appearances, condensation is a thermodynamically irreversible process (*42, 43*); the appearance of reversibility (Figure 1E-H) depends upon returning to a point beyond the lower or upper closure points of the hysteresis loop. The absence of a break in the curve, as would be expected from a concerted change in state of the entire system, is explained by heterogeneity of the material (discussed below). The chromosomal phase transition was apparent macroscopically by the sudden opacification of chromosomes in suspension upon addition of spermidine (Figure 4C).

Further evidence for a VPT came from the observation of phase coexistence — theoretically attainable within the zone bounded by binodal and spinodal lines — at low polyamine concentration (Figure 4D). Beginning from a phase-separated state, movement of the phase boundary was observed upon flowing solution favoring one of the two phases, resulting in the growth of that phase at the expense of the other (Figure 4E). *We conclude that chromosome condensation involves a volume phase transition of the chromosomal hydrogel*.

We note that a VPT is a reflection of the coil-globule transition of the polymer component of a gel (*44*). An intramolecular coil-globule transition has been observed for DNA in the presence of trivalent cations (*45, 46*). Therefore, the long-studied condensation of DNA and chromatin fragments manifests as a VPT when assembled into a network (Figure 4F).

## Macromolecule-driven condensation

The excluded volume effect is known to play a role in the compaction of naked DNA and of prokaryotic and eukaryotic genomes (*47–49*). The condensation of DNA by polyethylene glycol (PEG) requires sufficient counterions to screen repulsive electrostatic interactions in the condensed material. Precipitation of DNA by PEG is commonly performed in the presence of at least 0.5 M NaCl; in the presence of 0.2 M NaCl, DNA resists precipitation by PEG concentrations as high as 12% (*47*). In contrast, chromosomes were efficiently condensed by 6.5% PEG at a molar ionic strength below 0.01 (Figure 5A). The difference in behavior between DNA and chromosomes may be attributed to the histone proteins, which partially neutralize the charge of the DNA in chromatin and thereby diminish the strong repulsive forces that would otherwise oppose close packing in the condensed state. Whereas chromosome condensation was first order in polyamine concentration (Figure 3A), it was sixth order in PEG concentration (Figure 5A), indicative of a highly cooperative process. The dielectric constant of the solution was negligibly affected over the range of PEG concentrations tested (*50*), and therefore cannot explain the observed behavior. The mechanism of PEG-mediated condensation involves the excluded volume effect, reduction of water activity, and, to the extent that PEG is non-penetrating, osmotic pressure.

**Figure 5:**
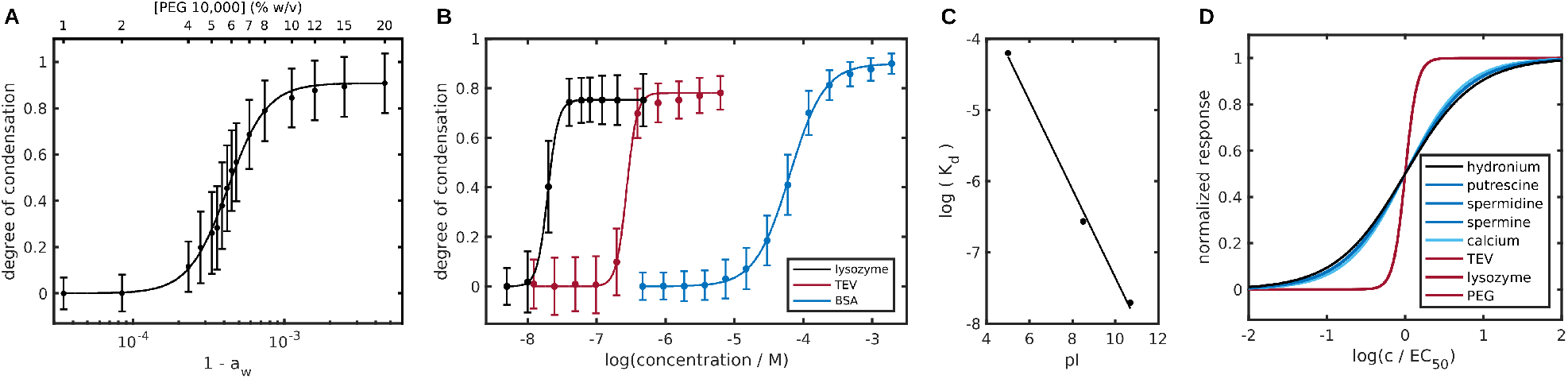
Molecular crowding and electrostatic bridging elicit chromosome condensation. (A) Swollen chromosomes were titrated with PEG 10,000 at concentrations ranging from 0 to 20% weight/volume. The normalized degree of condensation averaged over *n* = 372 chromosomes is plotted with respect to one minus the thermodynamic activity of water (lower abscissa) and the concentration (% w/v) of PEG 10,000 (upper abscissa). Note that the relation between water activity and PEG concentration is nonlinear. (B) Decondensed chromosomes were titrated with chicken egg lysozyme (black), catalytically inactive tobacco etch virus (TEV) protease (red), and bovine serum albumin (BSA; blue). The normalized degree of condensation was determined by averaging over *n* = 497, *n* = 551, and *n* = 772 chromosomes respectively. Error bars for (A) and (B) denote ±1*σ*. (C) Dependence of the apparent microscopic dissociation constant (*K*_d_) on protein isoelectric point (pI). The data are fit by a straight line with *r*^2^ = 0.995. (D) Comparison of response sensitivity by superposition of best-fit condensation profiles. The normalized degree of condensation is plotted with respect to logarithmic concentration, normalized by the EC_50_ of each agent.

Highly cooperative chromosome condensation (up to sixth-order dependence) was also caused by proteins, apparently nonspecifically, as it was induced by all proteins of molecular weights 10–100 kDa tested, including bovine serum albumin (BSA), chicken egg lysozyme, and catalytically inactive tobacco etch virus (TEV) protease (Figure 5B). The effect was modulated by charge neutralization, as basic proteins were considerably more potent than acidic ones. Differences in efficacy were explained by differences in the isoelectric points of the proteins (Figure 5C). Nevertheless, charge neutralization alone could not explain the effect, since acidic proteins — which augment fixed charge and thereby promote swelling — also induced condensation. Furthermore, the high degree of cooperativity compared with counterion-driven condensation (Figure 3A) indicated a role for factors beyond neutralization. Protein-mediated condensation was likely due to bridging between chromatin fibers. Bridging occurs through the interaction of charged elements of the fibers (for example negatively charged DNA phosphates and positively charged histone amino terminal tails) with sites of opposite charge on the proteins.

The sensitivity of the chromosomal response to environmental perturbation varied over a wide range (Figure 5D), and in a manner that depended upon the particular effect of a perturbation on chromatin. The response was gradual for perturbations that brought about a reduction in the ion osmotic pressure (black and blue curves in Figure 5D), whereas the response was highly cooperative for perturbations that acted through macromolecular crowding or fiber bridging (red curves in Figure 5D). Because chromatin, both mitotic and interphase, was condensed in almost any medium other than nearly pure water, *condensation may be regarded as the favored or “default” state of the chromosomal material in the cellular milieu, with its high concentrations of protein and multivalent cations*.

## Relevance to chromosome condensation-decondensation *in nucleo*

Swelling and deswelling of chromatin is not specific to the mitotic state (*51*). Nuclei exhibit similar behavior, swelling in low salt and collapsing in the presence of multivalent ions (Figure S2B). The nature of chromatin as an ionic hydrogel thus pertains in the interphase nucleus as well. In the interphase nucleus, chromatin is partitioned into so-called topologically associating domains, or “TADs” (*52, 53*), identified by an apparent enrichment of interactions within a domain compared to interactions with sites outside the domain. Detection of such interactions routinely involves the use of a chemical cross-linker (*54*). By examining the effect of chemical crosslinking on condensation behavior, we found that cross-linking preserves chromatin conformation only in the condensed state. Cross-linking of decondensed chromosomes had no apparent effect on their condensation-decondensation behavior (Figure 6A, top), whereas similar treatment of condensed chromosomes left them irreversibly condensed (Figure 6A, bottom). The average behavior for more than a hundred chromosomes (Figure 6B) was similar to that of nuclei (Figure 6C). The dependence of TAD detection on cross-linking (*54*) simply reflects the introduction of cross-links in condensed regions (Figures 6A and 6C, lower panels) but not decondensed ones (Figures 6A and 6C, upper panels), facilitated by differences in local chromatin density. TADs may be identified on this basis as condensed regions, consistent with previous studies on Drosophila polytene chromosomes (*55, 56*).

**Figure 6:**
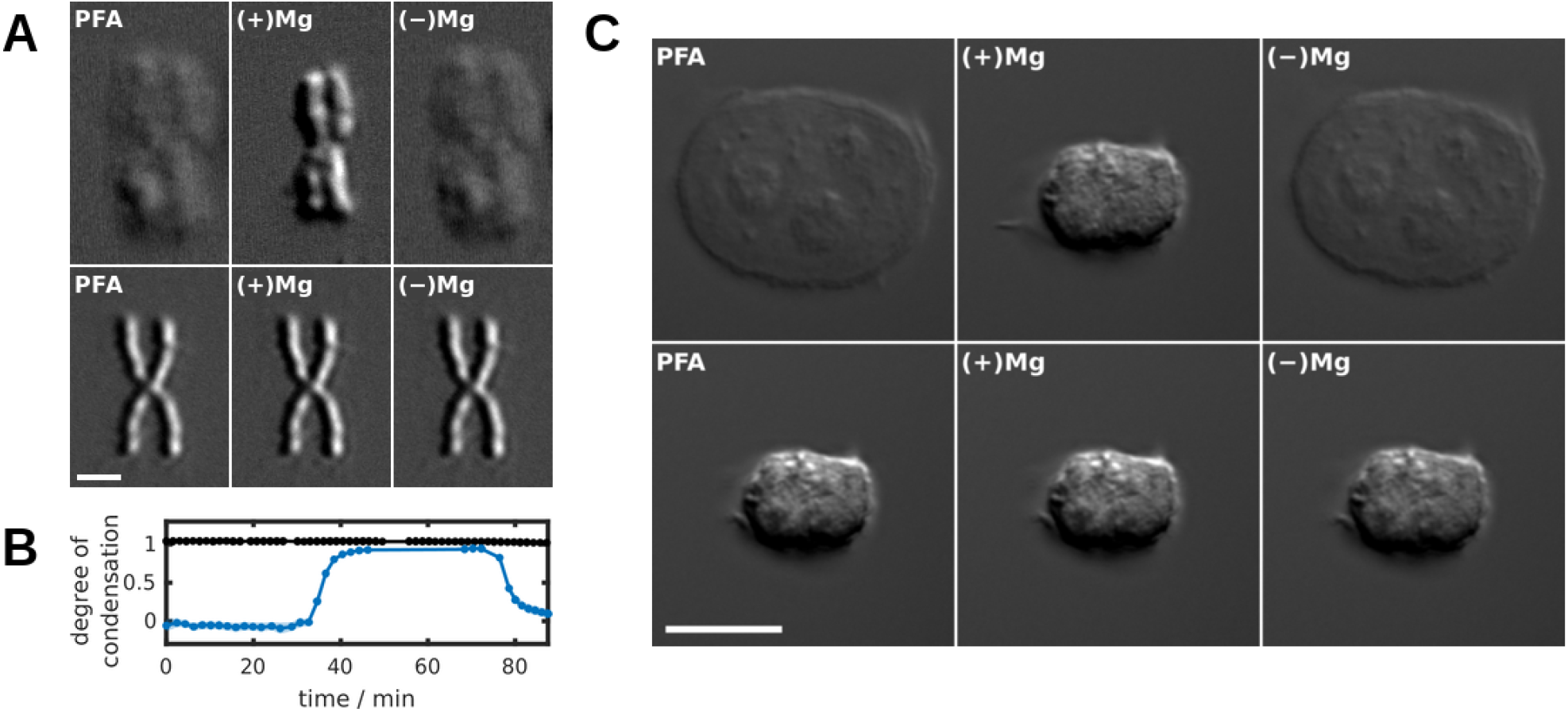
Relevance to chromosome condensation-decondensation *in nucleo*. (A) *Top*: Decondensed chromosomes cross-linked with 1% paraformaldehyde (PFA) in 5 mM HEPES (pH 7.5), 2 mM KCl (left) retained the ability to undergo condensation (middle) and decondensation (right). *Bottom*: Conversely, condensed chromosomes cross-linked with 1% PFA in 5 mM HEPES (pH 7.5), 2 mM KCl, 10 mM MgCl_2_ (left) lost the ability to decondense (right). Scale bar: 2 *µ*m. (B) Quantification of the behavior shown in panel (A) for chromosomes cross-linked in condensed (black) and decondensed (blue) states. Chromosomes were exposed to 1% PFA for thirty minutes. After washing to remove PFA, the chromosomes were immersed in condensing solution (5 mM HEPES [pH 7.5], 2 mM KCl, 10 mM MgCl_2_) followed by decondensing solution (5 mM HEPES [pH 7.5], 2 mM KCl). (C) *Top*: Decondensed nuclei cross-linked with 1% PFA in 5 mM HEPES (pH 7.5), 2 mM KCl (left) retained the ability to undergo condensation (middle) and decondensation (right). *Bottom*: Conversely, condensed nuclei cross-linked with 1% PFA in 5 mM HEPES (pH 7.5), 2 mM KCl, 10 mM MgCl_2_ (left) lost the ability to decondense (right). Scale bar: 10 *µ*m.

## Principles of mitotic chromosome structure

Mitotic chromosome structure derives from two organizing principles, a linear configuration of chromatin loops and a volume phase transition of the chromatin hydrogel. Chromosomal cylindricity and uniformity of width emerge simply from the minimization of surface area along a linear axis. The condensed state is favored, and largely maintained in interphase, by the conditions of the cellular milieu. Condensation by collapse of a hydrogel resolves the central paradox of the mitotic chromosome, the organization of a heterogeneous fiber in a regular structure. Collapse is compatible with local variation within an overall condensed structure.

Relaxation of the forces responsible for condensation is commonly thought to explain the uncoiling of chromosomal material in the interphase state. We find, however, that the condensed state is the “default” state, pertaining in interphase except in regions where, for purposes such as gene activity, decondensation is driven by forces or mechanisms that remain to be determined.

## Chromatin structural disorder

Hysteresis is a non-equilibrium phenomenon; despite being apparently smooth, hysteresis loops consist of a series of microscopically discrete, irreversible transitions or “jumps” (*57*). The precise shape of a hysteresis loop depends upon detailed material properties — steeper loops signify fewer jumps and more regular structure, while broader loops signify more jumps and less regular structure. If in the case of chromatin fibers the pattern of coiling or folding were relatively uniform, then so too would be the energy barrier to structural change, and transitions would occur over a narrow range of effector concentration. If instead the coiling or folding were variable, then the energy barrier to structural change would also vary, resulting in more gradual transitions. The breadth of the observed chromosome condensation-decondensation hysteresis loops (Figure 4A) therefore points to a significant degree of chromatin structural disorder. Chromatin evidently adopts heterogeneous and disordered conformations in both condensed and decondensed states.

## Consistency with prior evidence

The idea of chromosomes as gels is consistent with previous suggestions and experimental findings (*7, 16, 58*). The composition of chromosomes lends itself to gel formation, with an abundant polyelectrolyte, amenable to cross-linking by various agents, including multivalent cations (*59*), histone tails (*60*), and non-histone proteins (*61, 62*). Our work has placed the gel idea on a firm basis, going beyond conjecture and pertinent observations to a definitive experimental and theoretical identification of the chromosomal material as an ionic hydrogel. The concordance of our quantitative measurements with classical physicochemical theory, adapted for chromosomal material, led us to the discovery of the fundamental basis for chromosome condensation, collapse of the gel through a VPT.

Chromosome condensation-decondensation has been variously attributed to the formation and unraveling of hierarchical chromatin structure (*16,63,64*), the assembly and disassembly of multi-layered plate structures (*65*), and the formation and dissolution of a polymer melt (*66, 67*). The involvement of liquid-liquid phase separation has also recently been suggested (*68, 69*). A polymer melt is a neat, polymeric liquid above its glass transition or crystallization temperature, whereas a polymer gel is a “soft solid” comprising polymer and solvent (*70*). Unlike liquidliquid phase separation, mixing of polymer and solvent (embodied in our model by Equation 2) is just one of several processes determining the state of a gel. Elasticity — which chromosomes are known to possess (*71*) and which is embodied in our model by Equation 3 — is difficult to reconcile with a polymer melt or a phase-separated liquid mixture. Melts may respond elastically to a deformation over short timescales, but like other liquids, they ultimately yield to viscous flow. By contrast, gels possess rigidity, allowing them to maintain their shape under deformation. Elasticity explains why chromosomes do not dissolve under solution conditions (Figure 3A) which bring about the dissolution of chromatin condensates (*69*). In addition to mixing and elastic forces, the structure of polyelectrolyte gels is determined by an ion osmotic pressure (embodied in our model by Equation 4). This pressure explains the distention of chromosomes in low salt (Figure 3C). Structures that effectively retain fixed charge, including networks and semipermeable membranes (*24*), give rise to stable ion pressures and can assume a particular swollen state indefinitely under fixed conditions. In contrast, structures that permit the egress of fixed charge, including polymer melts and phase-separated liquid droplets, would be incompatible with the ion pressure observed. The surface tension of a liquid droplet, for instance, hinders but does not prevent the diffusion of its contents. The condensation profiles observed could not be fit by models lacking elastic or ion osmotic terms, indicating that simple binary mixing implied by liquid-liquid phase separation (*70*) cannot explain the observed behavior. Recent findings of liquid-liquid phase separation behavior in chromatin likely reflect the coil-globule transition of chromatin lacking persistent cross-links (top half of Figure 4F).

Condensation by collapse of a hydrogel is consistent with many observations, including stable organization with regional mobility, local heterogeneity of condensation state, morphological memory, cessation of DNA transactions during mitosis, chromosome cylindricity (in the presence of a scaffold), and Hi-C contact frequencies. The positional stability of a gel enables local structural variation; specific regions may be decondensed by targeted disruption of crosslinks, by local control of polymer properties such as charge density, and so forth. Inasmuch as hysteresis implies a form of morphological memory, local changes in chromatin condensation could persist even after removal of a stimulus, such as a local change in cross-linking or polymer properties. Condensation is fundamentally a syneresis (removal of water); the reduction in intrachromosomal water activity, and the barrier to diffusion posed by the high density of the collapsed state may explain the cessation of DNA transactions during mitosis. Finally, as mentioned, the cylindricity of mitotic chromosomes emerges as a basic property of an axially organized gel. An unscaffolded chromatin network model was previously proposed on the basis of the loss of chromosomal elasticity upon nuclease digestion (*7*). Such a model is difficult to reconcile with the linearity of chromosomes. In contrast, we observe a central filament by light microscopy which colocalizes with putative scaffold components and can be isolated as a discrete entity by nuclease digestion of chromosomes. The existence of a scaffold is also supported by a recent Hi-C analysis of metaphase chromosomes (*5*).

Previous Hi-C analysis of mitotic chromosomes demonstrated a loss of chromosome compartments and topologically associating domains (TADs) (*72*). Relatively uniform Hi-C contact frequencies have been observed for metaphase chromosomes, with a slow decline of contact frequency out to 10 Mb (*72*), and within TADs of the interphase nucleus (*52, 53*). The apparent contact of chromatin at one location in a region with chromatin at every other location may be explained by an ensemble average over many collapsed hydrogels. In interphase, the chromatin hydrogel is partitioned by region-specific decondensation into TADs, which are maintained by structural proteins such as CTCF. Upon entry to mitosis, the removal of these boundary proteins (*73*) may trigger the disappearance of TADs and chromosome-wide hydrogel collapse. Collapsed chromatin hydrogels are susceptible to cross-linking, whereas swollen ones are not (Figure 6), pointing to a relationship between TADs and the collapsed gel phase.

## Relevance to physiologic condensation

Our demonstration of the hydrogel nature of the chromosome relied on its propensity to swell in low-salt solutions. Nevertheless, the generality of our conclusions does not depend on particular solution conditions. Indeed, a gel will remain a gel so long as its cross-links, be they static or dynamic, persist. Raising the ionic strength to physiologic levels will not alter the nature of the chromosome as a gel, though it will increase the degree of condensation. Indeed, at physiologic ionic strength (log *c ≈ −* 0.8 in Figure 3I), the chromosome is largely (though not fully) condensed. Likewise, *in vivo*, most chromatin is condensed most of the time. Only limited regions are subjected to decondensation, for purposes of gene activity and so forth, as dictated by the metabolic needs of the cell. The preponderance of condensed chromatin *in vivo* is explained by our observation that condensation is the natural or “default” state of chromatin in physiologic solution. How then is regioselective decondensation of the chromosomal gel achieved? The most probable explanation is through local modification of network properties, including charge and cross-link density, mediated by site-specific histone-modifying and chromatin-remodeling activities. As noted, most negative charge resides in the DNA phosphates, while most positive charge resides in the histones, especially in their amino-terminal tails. The theory of polyelectrolyte gels rationalizes how post-translational modifications affecting the charges of the tails can influence condensation. All histone post-translational modifications will affect mixing thermodynamics through effects on *χ* (Equation 2) that are difficult to predict quantitatively, and some, like lysine acetylation, will also affect the charge density of the gel (*z*_2_ in Equation 5). Various processes will affect the density of intrachromosomal cross-linking (*N_x_* in Equation 3), including histone posttranslational modifications (e.g., H4K16ac (*74*)), changes in concentrations of polyvalent ions, and topological interactions (e.g., mediated by various SMC complexes (*62*)). Among these effects, variation of *χ* has the most profound effect on chromosome volume at physiologic ionic strength, where it is predicted to induce a VPT (Figure S7).

The cellular mediators of mitotic chromosome condensation remain to be established. The level of histone acetylation is significantly reduced around the time of entry to mitosis (*75, 76*). Preventing this loss by inhibition of histone deacetylase activity prevents chromosome condensation (*77*). The basis for the effect appears to reside in the post-translational modification state of the histones (*78*). These findings are readily explained by our model (Figure S7): histone deacetylation effectively reduces the negative charge density of chromatin (i.e., reduces the magnitude of *z*_2_) and is predicted to diminish chromatin solubility through effects on *χ* favoring demixing. In addition to the effects of histone post-translational modifications, the widespread condensation seen during mitosis is likely promoted by changes in the cellular milieu, such as by increases in the concentration of magnesium (*79*) and polyamines (*80*). The bound pool of polyamines undergoes a dramatic, transient rise at the time of cell division (*80*), which may be sufficient to induce a volume phase transition. Indeed, the degree of condensation of chromosomes in solutions of physiologic ionic strength (about three quarters, using the convention adopted herein) is similar to that at which condensation reactions become thermodynamically irreversible (compare Figures 3I and 4A), suggesting that chromosomes are poised to exploit the properties of the adjacent volume phase transition.

A transition to a more condensed phase entails attractive interactions, which form noncovalent cross-links in a hydrogel. Our findings demonstrate the importance of charge-charge interactions in the condensation of a chromatin hydrogel. Additional interactions of the tails and other proteins may modulate condensation as well. The identification and roles of such interactions in the VPT and in the reversal of the VPT for decondensation in interphase remain to be determined. Other outstanding questions include the assembly of the central filament, or scaffold, the trajectories of chromatin fibers in the condensed state, and the relationship to gene activity.

## Acknowledgments

We thank Peter Geiduschek for reviewing this manuscript, Snezana Djordjevic for the generous gift of TEV protease expression plasmids, Geoffrey Wahl and Teru Kanda for generously providing the H2B-GFP-expressing HeLa cell line, and Jon Mulholland and the Stanford Cell Sciences Imaging Facility for microscope access and training.

## Contribution

All authors contributed to study design and writing of the manuscript. A.J.B and P-J M. performed the experiments. A.J.B performed the computational analysis.

## Data and materials availability

Raw data will be made available upon request.

## Supplementary Materials

### Materials and Methods

#### Chromosome purification

HeLa S3 suspension cells were grown in Minimal Essential Medium supplemented with 10% newborn calf serum, and adherent H2B-GFP HeLa cells (*81*) were grown in Dulbecco’s Modified Eagle Medium containing 7.5% fetal bovine serum. Cells were grown at 37 °C in a 5% CO_2_ atmosphere, and were arrested in mitosis by treatment with 100 ng/mL colcemid for 16 h (mitotic index *∼*0.6). Mitotic chromosomes were purified according to a protocol derived from the polyamine-based purification method of Lewis and Laemmli (*2*). Specifically, arrested cells were harvested by centrifuging at 1,000 g for 10’ at room temperature and then resuspended in one tenth of the culture volume of a solution consisting of 7.5 mM Tris-HCl (pH 7.5), 40 mM KCl, 1 mM EDTA, 0.375 mM spermidine, 0.15 mM spermine, 1% thiodiethanol (v/v), and protease inhibitors (1 mM benzamidine, 100 µM leupeptin, 10 µM pepstatin A, 1 mM phenylmethylsulfonyl fluoride). Cells were permitted to swell at room temperature for 10’, and were subsequently pelleted (600 g for 6’), after which the supernatant was discarded. The cell pellet was resuspended in 10 mL of lysis buffer (twice concentrated swelling buffer supplemented with 0.1% [w/v] digitonin), and lysis was achieved by 12 strokes of a loose-fitting pestle. The lysate was loaded onto a 3 x 10 mL discontinuous gradient of sucrose (15%, 50%, 80% w/v) in a buffer consisting of 5 mM Tris-HCl (pH 7.5), 2 mM KCl, 2 mM EDTA, 0.375 mM spermidine, 1% thiodiethanol (v/v), 0.1% lauryldimethylamine oxide (w/v), and protease inhibitors, and then spun at 1,500 g for 45’. The chromosomes at the 50-80% interface were diluted to 40 mL with 88% Percoll in the previous buffer supplemented with spermidine and spermine to final concentrations of 2 mM and 0.8 mM, respectively, and centrifuged at 45,000 g for 20’. The chromosomes, which formed a diffuse band near the bottom of the tube, were recovered and washed with a solution consisting of 5 mM Tris-HCl (pH 7.5), 2 mM KCl, and 2 mM EDTA, pelleted at 3,000 g for 10’, and finally resuspended in a solution comprising 5 mM Tris-HCl (pH 7.5), 2 mM KCl, and 0.01% (w/v) sodium azide.

#### Nuclei purification

HeLa S3 cells were washed with PBS and resuspended in ice-cold hypotonic buffer (10 mM HEPES-KOH [pH 7.5], 25 mM KCl, 2 mM MgCl_2_, 1 mM dithiothreitol, and protease inhibitors), centrifuged, and resuspended in ten volumes of the previous buffer. The cells were left on ice for 1 h and subsequently Dounce-homogenized using a loose-fitting pestle. Sucrose was added to the lysate to a final concentration of 250 mM. The nuclei were pelleted (1,000 g, 10’), washed once more in ten volumes of a similar buffer (10 mM HEPESKOH [pH 7.5], 25 mM KCl, 2 mM MgCl_2_, 250 mM sucrose), resuspended in that buffer to a numerical concentration of 10^6^ nuclei per microliter, and stored under liquid nitrogen. Thawed nuclear suspension was diluted 10-fold in 10 mM Tris (pH 7.5), 25 mM KCl, 2 mM MgCl_2_, and settled onto mPEG-AEPTES-derivatized coverslips for perfusion studies.

#### Coverslip derivatization

Coverslips of 40 mm diameter and 170 *±* 5 µm thickness (Paul Marienfeld, GmbH) were washed in a 2% (v/v) alkaline detergent solution (Mucasol, Schülke) for 30’ in a bath sonicator. They were subsequently washed five times in 18.2 MΩ *·* cm (Milli-Q) water and sonicated for an additional 5’ in Milli-Q water before being immersed in spectrophotometry-grade acetone (Photrex, J.T. Baker) and allowed to dry at room temperature. A silanol layer was produced by exposing the coverslips to air plasma (Harrick Plasma PDC-32G, set to high power) for 5’ and allowed to sit for 15’, following which derivatization commenced by dipping in acetone and then immersing for 10 s in acetone supplemented with 2% (v/v) 3-(2-aminoethylamino)propyltriethoxysilane (AEPTES, Gelest, Inc.). Following two additional acetone washes, the coverslips were dried on a hot plate set to 170°C for 10’ and bath-sonicated in absolute ethanol for 10’. After washing extensively with water, they were finally dipped in acetone and allowed to dry before being stored at room temperature under vacuum. A layer of PEG was applied by diluting mPEG-succinimidyl valerate (mPEG-SVA, MW 5,000, Laysan Bio, Inc.) to 0.1 mg/mL in 100 mM potassium perborate (pH 8.4), and laying AEPTES-derivatized coverslips on a 100 µL drop of that solution for 2 h. Coverslips were then rinsed five times with Milli-Q water, and allowed to dry before being stored under vacuum.

#### Widefield microscopy

Chromosomes (0.08 A_260_ units) were adsorbed to amine-functionalized coverslips, mounted in an FCS2 perfusion chamber (Bioptechs), allowed to settle for 1 h at room temperature, and washed with 5 mM Tris-HCl (pH 7.5) and 2 mM KCl delivered from a Harvard Apparatus 11 Plus syringe pump at a flow rate of 50 µL/min. The chamber was mounted in a K-stage adapter (160 x 110 mm) and loaded on a Nikon Eclipse-TI inverted microscope equipped with a Plan-Apo-TIRF, 100x, NA 1.49, oil-immersion objective. Micrographs were collected on an Andor Neo sCMOS camera with a 2560×2160-pixel sensor (6.5 µm pixel size). For each time point, a z-stack comprising 20 to 40 Nyquist-sampled slices was collected using a piezoelectric z-drive. Buffer exchange was performed at flow rates not exceeding 500 µL/min and adjustments to the flow rate were graduated to avoid deleterious shear gradients. We note that our experimental system allows for precise specification of the free concentration of ligand ([L]_free_), which is convenient for the study of associative processes. All solutions were made using a base of 5 mM Tris-HCl (pH 7.5) and 2 mM KCl, with the exception of those employed for the pH and ionic strength titrations. For the pH titration, Tris-HCl was replaced by 5 mM concentrations of the following buffers: glycine-HCl (for the pH range 2.5–3.5), potassium acetate (pH 3.75–5.75), PIPES-KOH (pH 6.0–7.0), and glycine-KOH (for pH greater than 8.5). For the ionic strength titration, solutions of KCl were prepared in argon-saturated water to exclude atmospheric carbon dioxide. PEG concentrations (Figure 5A) were expressed in terms of the corresponding thermodynamic activity of water using published measurements of density and activity (*82–84*). Lyophilized proteins (BSA [Calbiochem], hen egg white lysozyme [SigmaMillipore]) were diluted in 5 mM Tris (pH 7.5), 2 mM KCl. TEV protease (C151A mutant), lysozyme, and BSA solutions were dialyzed overnight against 5 mM Tris, 2 mM KCl, pH 7.5 to remove any trace of salt present in excess in the lyophilizate.

#### Condensation profile determination

Determination of condensation profiles from DIC micrographs was accomplished in multiple steps. Single chromosomes were windowed within fields of chromosomes, affording regions of interest (“ROIs”). The focal plane for each ROI z-stack was determined using the variance of Tenenbaum’s gradient (*85*), which we found to be among the most robust functions for focusing DIC z-stacks from a library of focus measures (*86*). ROI focal planes were then subjected to morphological determination of chromosome area using the following procedure: firstly, micrographs were low-pass filtered by convolution with a Gaussian kernel; secondly, contrast was enhanced by contrast-limited adaptive histogram equalization (*87*); thirdly, edge detection was performed using the Canny algorithm (*88*); fourthly, short, perimeter-adjacent edges were pruned with an efficiency related to the proximity thereof to the ROI perimeter; fifthly, the image was closed with a disk-shaped structuring element and local minima eliminated by a flood-fill operation; sixthly, the image was opened with a smaller structuring element than the one used for closing, the purpose of which was to prune spurs; seventhly, area opening was performed to remove elements with subthreshold connectivity; eighthly, the convex hull of the segmented foreground was computed; and finally, the convex hull was smoothed with reference to the underlying micrograph by an active contour algorithm (*89*). The areas computed for each chromosome were internally normalized with respect to the areas computed from two calibration points collected for each data set, namely 0 and 375 µM spermidine in 5 mM Tris-HCl (pH 7.5), 2 mM KCl. By this normalization procedure, the degree of condensation was assigned to be 0 or 1 in the presence of 0 or 375 µM spermidine, respectively; this measurement calibration allowed for the comparison of condensation profiles from different experiments. We note that this definition of the degree of condensation is related nonlinearly to density. Areas computed by this procedure agreed well with areas determined by manually delineating chromosome contours (Figure S3). By means of nonlinear least-squares regression, the normalized areas could be fit by the Hill equation, yielding estimates of the apparent dissociation constant and the Hill coefficient. Error bars displayed in plots of condensation profiles denote *±*1*σ* (i.e., one standard deviation in either direction, centered about the mean value), and logarithms noted in such plots are of decimal base. The charge at pH 7.5 of methylamine, putrescine, spermidine, and spermine were computed using published pKa values for the amine moieties (*90, 91*).

#### Chromosome volume determination

For the determination of chromosome volume, coverslips were further passivated by coating with bovine serum albumin (BSA) to weaken surface contacts that may constrain the accessible range of chromosome volume. Z-stacks of chromosomes were collected on a Zeiss LSM880 laser scanning confocal microscope equipped with an AiryScan detector while perfusing solutions containing 10 nM Sytox Green, well below the concentration observed to induce condensation (Figure 2A). Excitation light (488 nm) was provided by an argon laser directed through a 63*×*, NA 1.4 PlanApochromat oil-immersion objective lens, and rastered across the sample at a zoom setting of 1.8. Emitted light was collected at twice the Nyquist frequency (AiryScan super-resolution mode) by a 32-element gallium arsenide phosphide compound detector placed in the conjugate plane, and deconvolved in Zen software (Zeiss) using a Wiener filter setting of 6.0. The AiryScan detector was calibrated using 0.1 µm TetraSpeck fluorescent beads (Life Technologies). Two approaches were then used to measure chromosomal volume from deconvolved z-stacks; the first method involved segmentation by an Otsu threshold, and the second refined the Otsu mask by applying the Chan-Vese active contour algorithm with a mild contraction bias. Volumes computed by the first method were slightly larger than those by the latter, but the volumetric ratio (the measurement of interest), was similar for both.

### Supplementary Text

We adapted a classical mean-field theory of gel swelling (*24*) for the prediction of chromosome swelling profiles. In addition to the assumption of a weak-screening limit, we further assumed additive independence of the contributions to the system’s free energy, negligible contribution of mobile ions to the system’s volume, and inter-phase activity coefficient ratios of unity. These approximations are justified under dilute conditions, for which the predictive accuracy of the model is expected to be greatest. We also ignored dependence of the Flory-Huggins parameter on ionic strength as well as any effects related to surface adsorption (i.e., the swelling process was regarded as unconstrained). The equilibrium state minimizes the free energy of the system, or, equivalently, results in zero net pressure, which is expressed by

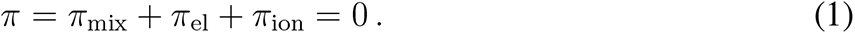

### The mixing term is modeled by Flory-Huggins solution theory, giving

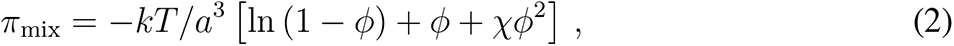

where *k* is the Boltzmann constant, *T* is the absolute temperature, *a*^3^ is the volume of an element of the lattice in terms of which the theory is formulated, *φ* is the polymer volume fraction (the volume fraction of solvent being 1 *− φ*), and *χ* is the Flory-Huggins parameter, which characterizes the energetics of the mixing process. The Flory-Huggins parameter is commonly expanded to first order in the polymer volume fraction (*χ* = *χ*_0_ + *χ*_1_*φ*), a practice that was adopted herein.

To account for finite chain extensibility, the elastic term was taken to be that for a network of Langevin chains (*92, 93*):

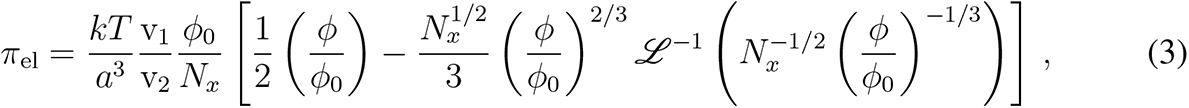

where *L ^−^*^1^ denotes the inverse Langevin function, v_1_ the molar volume of solvent, v_2_ the molar volume of a statistical chain segment, *N_x_* the degree of polymerization (i.e., the number of chain segments between cross-linking points), and *φ*_0_ the volume fraction of the gel in its reference state, in which chains assume ideal conformations.

### The ion osmotic pressure was approximated by the van’t Hoff formula

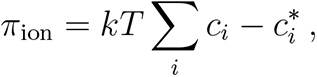

where *c_i_* and 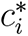 denote the concentrations of the *i*^th^ ionic species in the gel phase and the reservoir, respectively. The external concentrations are subject to experimental control and the internal concentrations are determined by Donnan equilibrium, which has a long history of application to the problem of polyelectrolyte gel swelling (*24,25,94–96*). Acid-dissociation and counterion-association reactions involving the gel’s ionizable groups were taken into account when calculating the fixed charge of the gel.

With these considerations in mind, the ion osmotic pressure and the electroneutrality constraint can be formulated in terms of the Donnan ratio *λ* in a manner that is readily extensible to arbitrary solution contexts. The ion osmotic pressure is described by

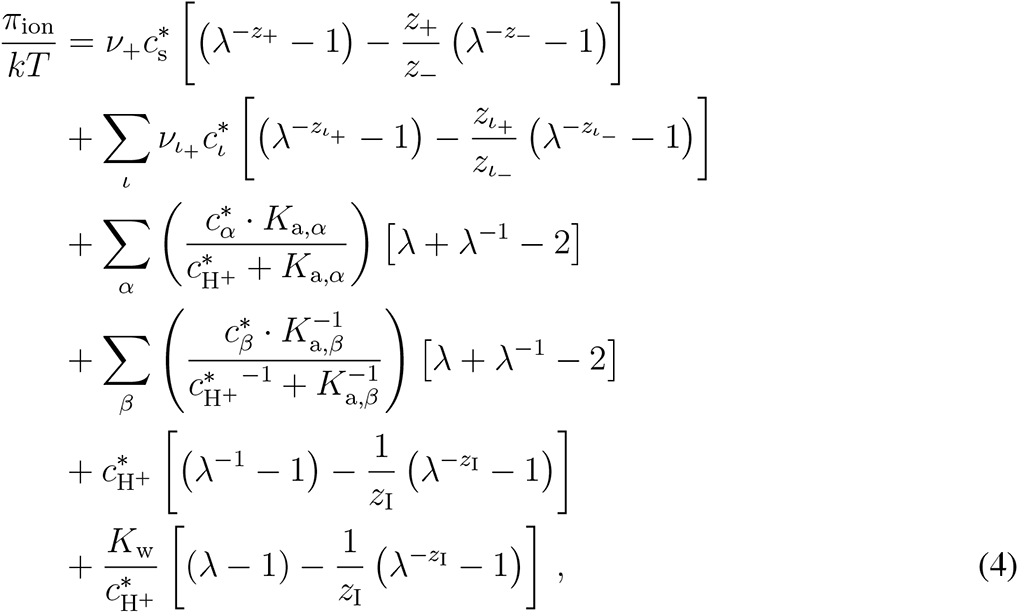

while the electroneutrality equation is given by

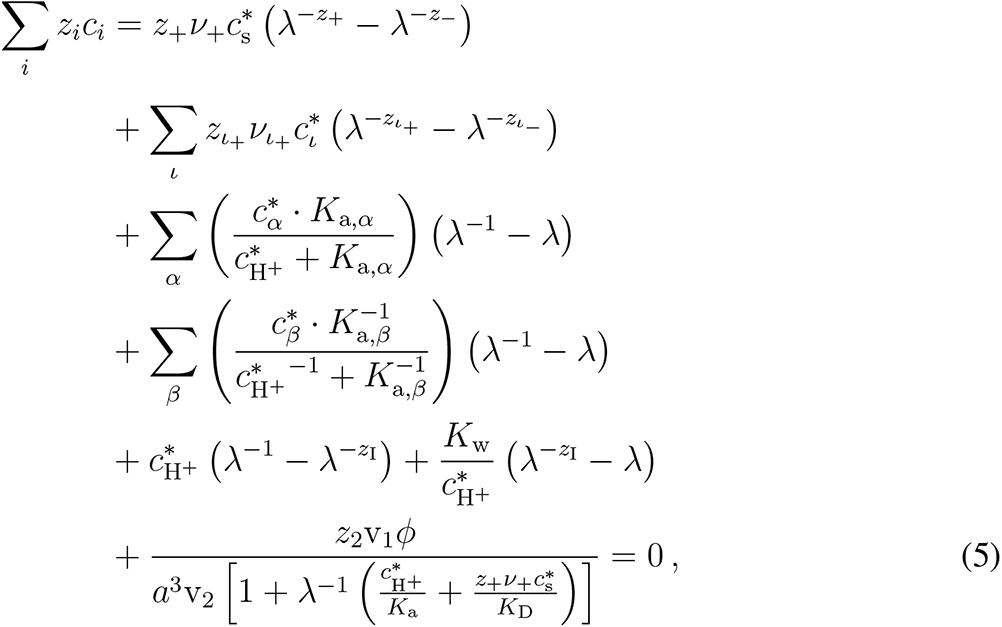

where the reservoir concentration of the titrated salt (composed of *ν*_+_ cations of valence *z*_+_ and *ν_−_* anions of valence *z_−_* per formula unit) is denoted by 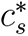. Background solutes are included in the summations over ionic compounds apart from the titrated salt (index *ι*), acids (index *α*), and bases (index *β*). Ionic compounds comprise cations of valence *z_ι_*_+_ and anions of valence *z_ι−_* represented in formula units with stoichiometric coefficients of *ν_ι_*_+_ and *ν_ι−_*. Monoprotic buffers with neutral acidic forms (having acid dissociation constants *K*_a_*_,α_*) or neutral basic forms (having conjugate acid dissociation constants *K*_a_*_,β_*) are present at concentrations of 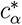 and 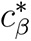, respectively. Hydronium and hydroxide are present at concentration of 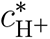 and 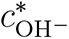 (which are linked according to *K*_w_, the autoionization constant of water), and the pH is set by an appropriate quantity of strong acid or base whose counterion bears an algebraic charge of *z*_I_. Network strands consist of monomers bearing a mean charge of *z*_2_ elementary units upon full dissociation. The association reactions of ionized groups with protons and counterions are governed by an acid dissociation constant *K*_a_ and a counterion dissociation constant *K*_D_, respectively.

The equation of state for the gel is therefore determined by simultaneous nonlinear equations, the numerical solution of which yields theoretical swelling profiles such as those depicted in Figures 3C and 3I.

Experimental data were fit to predictions from Equations 1–5 by a Monte-Carlo-guided random walk over the parameter space of the model. Free parameters included *φ*_0_, *N_x_*, v_2_, *z*_2_, *χ*_0_, *chi*_1_, *K*_a_, and *K*_D_, the latter of which was parameterized as a linear function of the counterion charge (*K*_D_ = *b*_0_ +*b*_1_*z*_+_). Data used for model refinement included the condensation profiles obtained for methylamine, putrescine, spermine, and KCl. The data for spermidine were omitted during model refinement. Mean chi-squared values (*χ*^2^*/n*) were computed according

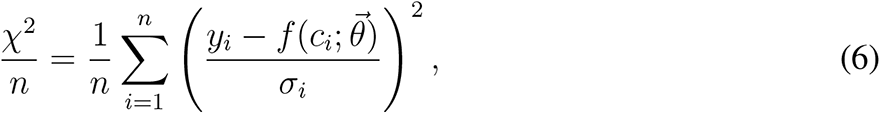

where *y_i_* is the observed degree of condensation at concentration *c_i_* with associated measurement error *σ_i_*, and 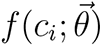 is the predicted degree of condensation at concentration *c_i_* under parameterization 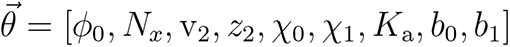.

## Supplementary Figures

**Figure S1:**
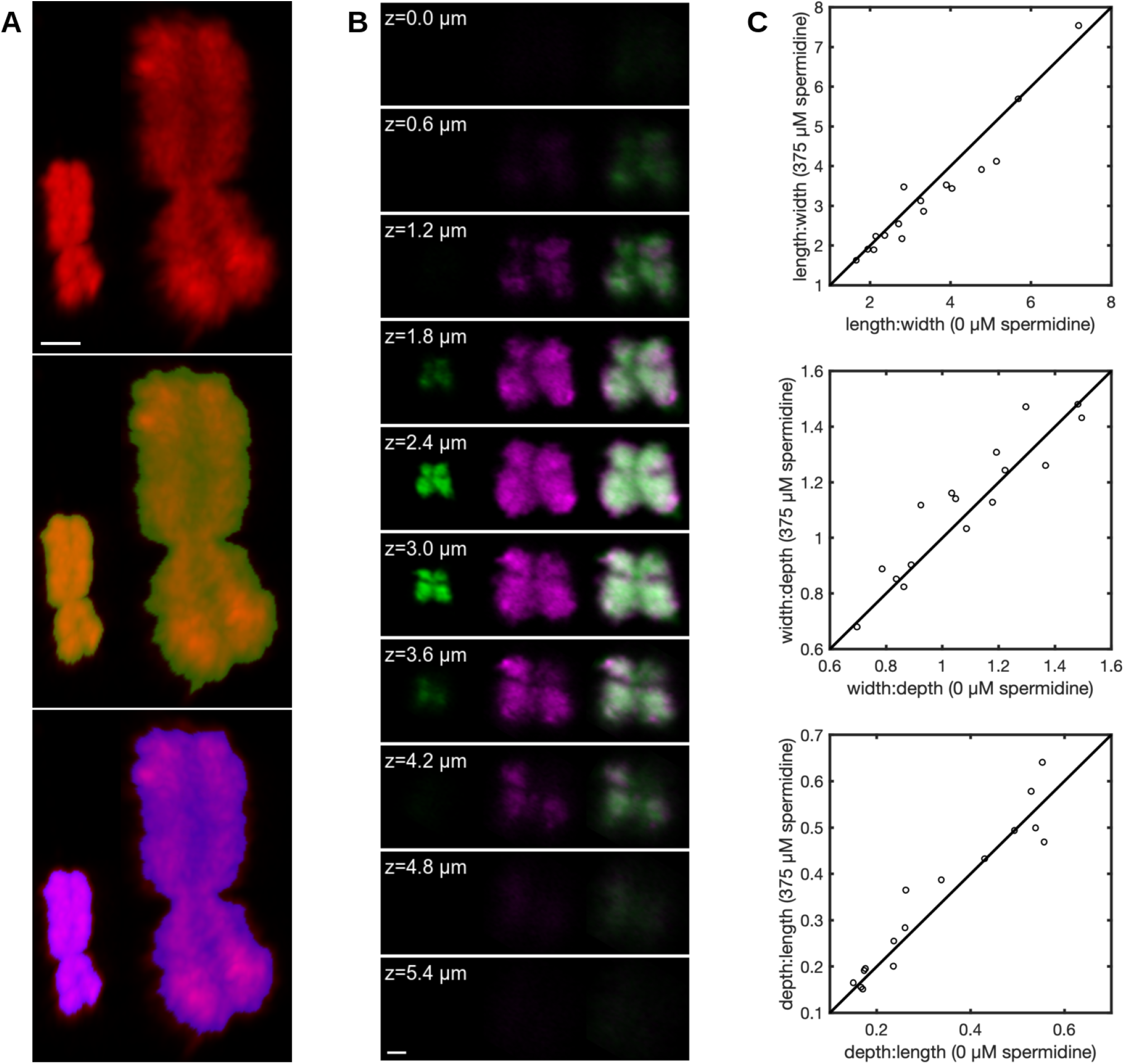
Three-dimensional analysis of chromosome morphology and condensation behavior. (A) Estimation of chromosome volume from z-stacks of Sytox Green-stained chromosomes (top; cf. Figure 1B) acquired on a laser-scanning confocal microscope (Zeiss LSM880) in the presence or absence of 375 µM spermidine. Chromosome volume was determined by means of an Otsu threshold alone (green mask, center) or in combination with an energy-minimizing active contour algorithm with a modest contraction bias (blue mask, bottom). The mean volume ratio between swollen and deswollen states for *n* = 54 chromosomes was 9.15 ± 0.34-fold by the former method and 9.95 ± 0.43-fold by the latter (mean ± S.E.M.). For visualization, Sytox Green was supplied at a concentration of 10 nM, significantly below levels observed to induce condensation (Figure 2A). Scale bar, 1 µm. (B) Optical sections (rows) of a chromosome in condensed (left third of each panel; green) and decondensed (middle third of each panel; magenta) states. The condensed chromosome was expanded isotropically *in silico* by the linear scaling factor (*V_d_/V_c_*)^1^*^/^*^3^, where *V_d_* and *V_c_* represent the volumes of the chromosome in decondensed (magenta) and condensed (green) states. The computationally expanded image of the condensed chromosome (right third of each panel; green) was brought into superposition with the decondensed state (right panels; magenta) by gradient-based optimization of the mean squared error of the three-dimensional images. A Pearson correlation coefficient was computed by pairwise comparison of pixel intensities throughout the superimposed three-dimensional volumes, shown in sections (right third of each panel), after normalization of intensities in each volume to the range [0,1]. The correlation coefficient for the chromosome shown here is 0.952. The spacing of the sections displayed is 0.6 µm; the scale bar represents 1 µm. (C) Immobilized chromosomes were imaged in two states of condensation obtained by equilibration with solutions containing 5 mM Tris (pH 7.5), 2 mM KCl, and either 0 µM or 375 µM spermidine. Image stacks for *n* = 16 chromosomes were subjected to measurements of chromosome length, width, and depth. Aspect ratios were computed along cyclically permuted dimensions. Points correspond to the ratio of the extent of the chromosomes along two spatial dimensions, as indicated by the axis labels, and lines describes the isotropic condition (*y* = *x*). Clustering of the data points about the line *y* = *x* demonstrates the isotropic nature of condensation. *Top*: length:width ratios at 0 µM and 375 µM spermidine. *Middle*: width:depth ratios at 0 µM and 375 µM spermidine. *Bottom*: depth:length ratios at 0 µM and 375 µM spermidine.

**Figure S2:**
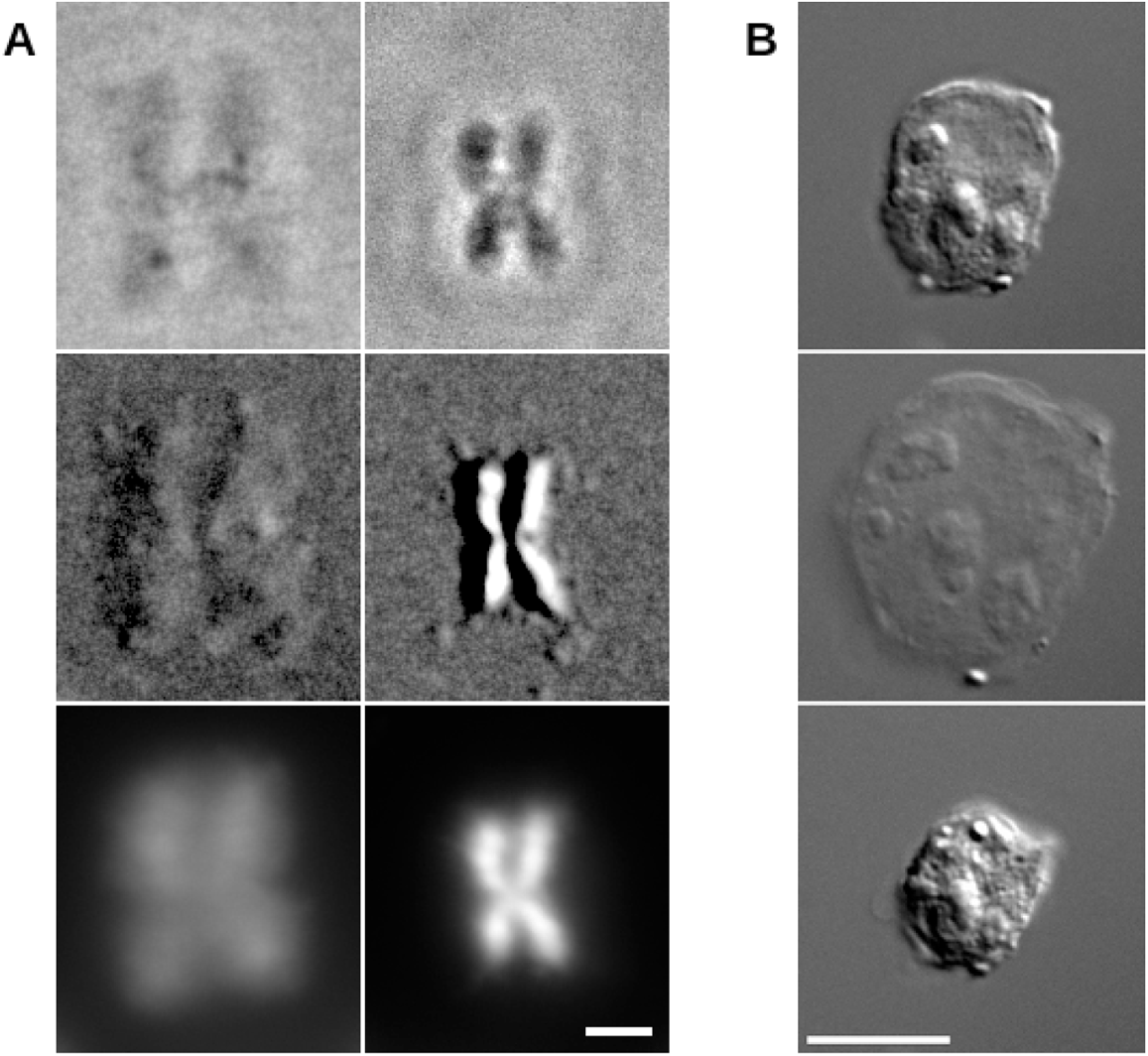
Swelling behavior of chromosomes and nuclei. (A) Correspondence between chromosome swelling/deswelling and chromatin decondensation/recondensation. A chromosome adsorbed to an amine-functionalized coverslip was exposed to spermidine concentrations of 0 µM (left) and 375 µM (right), corresponding to decondensed and condensed states of chromatin, respectively, and imaged by Zernike phase contrast (top), DIC (middle), and H2B-GFP fluorescence (bottom). The distribution of chromatin, represented by H2B-GFP fluorescence, mirrors changes in the specimen’s optical path length (phase contrast) or a spatial derivative thereof (DIC). Scale bar, 2 µm. (B) Nuclei swell and deswell under similar conditions to those for chromosomes. Purified HeLa cell nuclei were deposited on an amine-functionalized surface and imaged using DIC optics. Initially, the nuclei were placed in a solution consisting of 10 mM Tris-HCl (pH 7.5), 25 mM KCl, and 2 mM MgCl_2_ (top). They were swollen by flowing 5 mM Tris-HCl (pH 7.5) with 2 mM KCl (middle), and then deswollen by flowing the same buffer supplemented with 375 µM spermidine (bottom). Scale bar, 10 µm.

**Figure S3:**
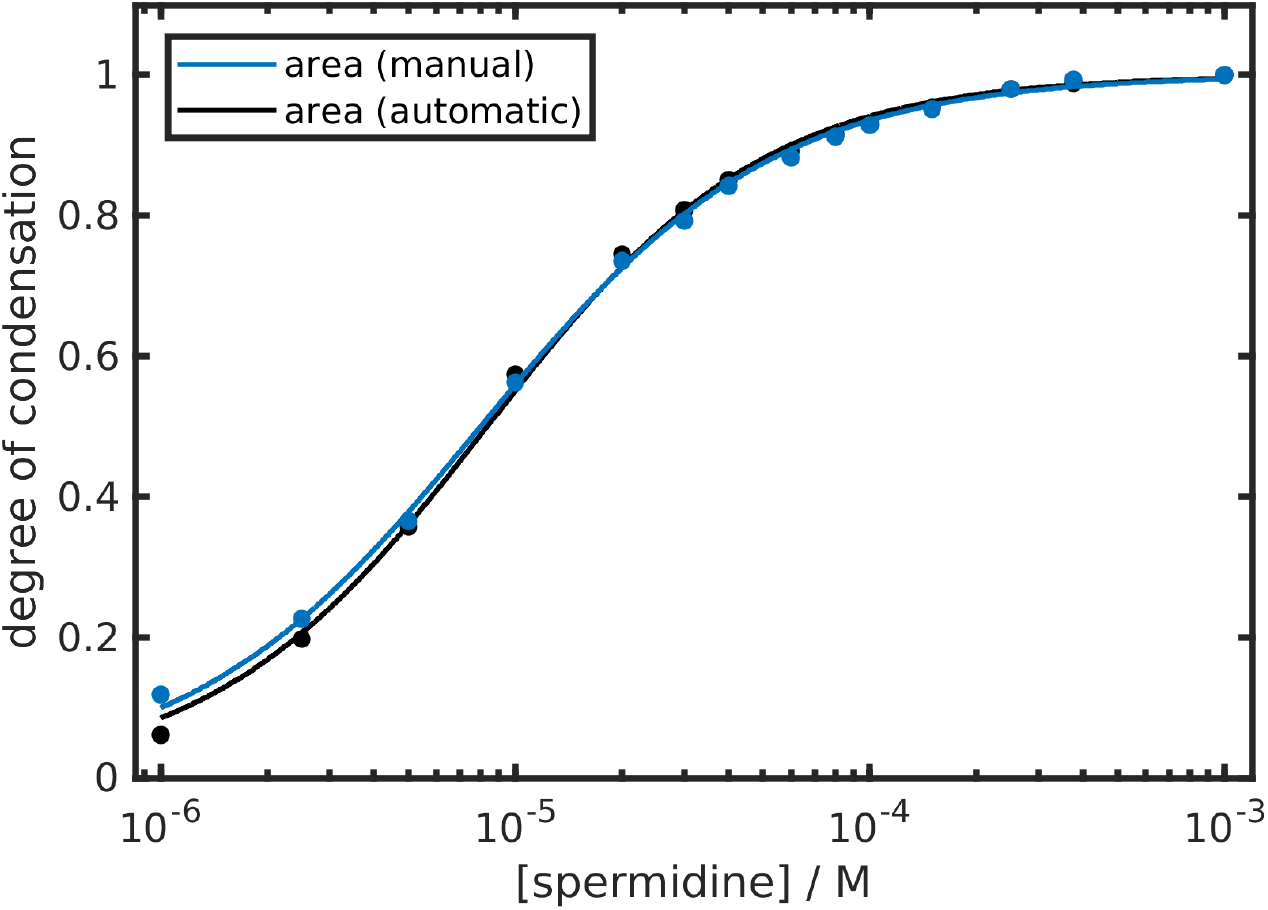
Validation of computational analysis of chromosome morphology. To validate the segmentation routine, images of chromosomes titrated with spermidine were segmented manually (black) and automatically (blue). The results of automatic segmentation shown here are also displayed as the black curve in Figure 2B.

**Figure S4:**
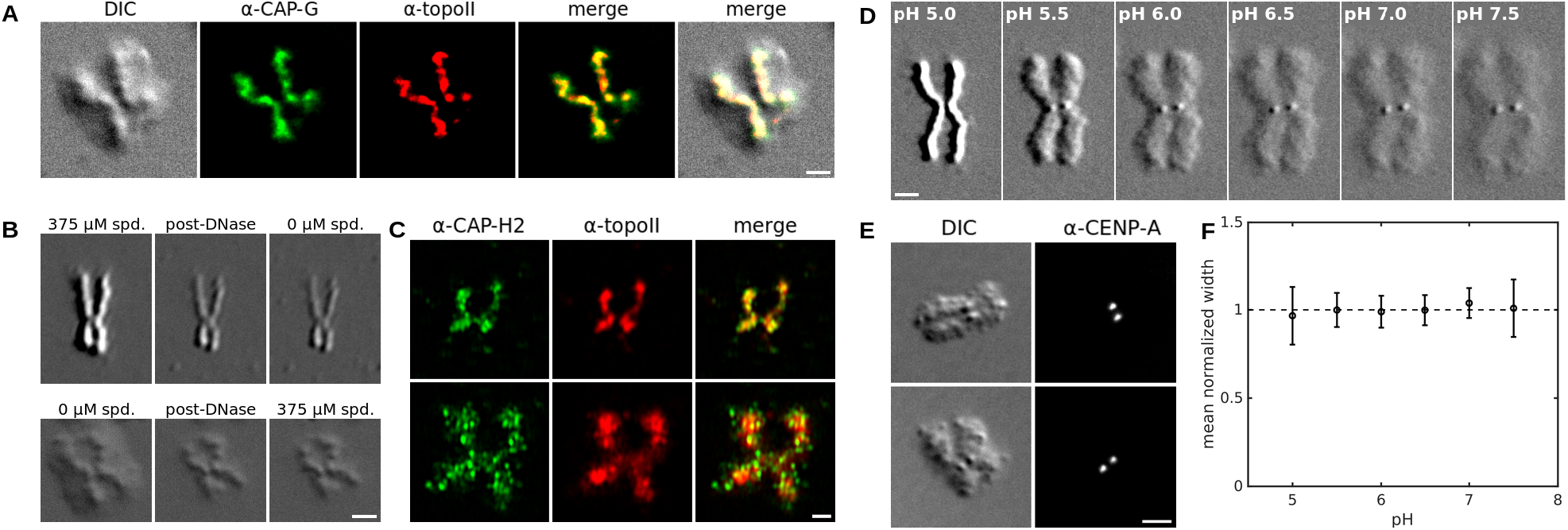
Direct observation of scaffold and kinetochores by chromosome decondensation. (A) Differential interference contrast image of a decondensed chromosome (left). Immunofluorescence (IF) with anti-hCAP-G (stained with Alexa Fluor 488 [green]) and anti-hTOP2A (topoisomerase II*α*, coupled to Alexa Fluor 647 [red]) shows colocalization of putative scaffold components with the central filament revealed by decondensation (three-way merge in right panel). Scale bar, 2 µm. The DIC and merged images are reproduced in Figure 3G. (B) The central filament is resistant to DNase digestion when examined in a condensed (upper panels) or in a decondensed (lower panels) state. The initial (undigested) state is shown in the left panels. The filaments resulting from digestion (middle panels) remain unchanged (right panels) when the condensing activity of the milieu is adjusted (from high to low [top] or *vice versa* [bottom]). (C) The scaffold contracts in the presence (375 µM, upper panels) and expands in the absence (0 µM, lower panels) of spermidine. (D) Kinetochores, recognizable as metacentric dots, are made visible by decondensation; they do not swell. (E) Identification of the metacentric dots as kinetochores by comparison of DIC and IF images using anti-CENP-A antibody coupled to Alexa Fluor 647. Two examples are shown. (F) The normalized mean diameter of 16 kinetochores is plotted as a function of the solution pH: kinetochore size remains constant during chromosome decondensation.

**Figure S5:**
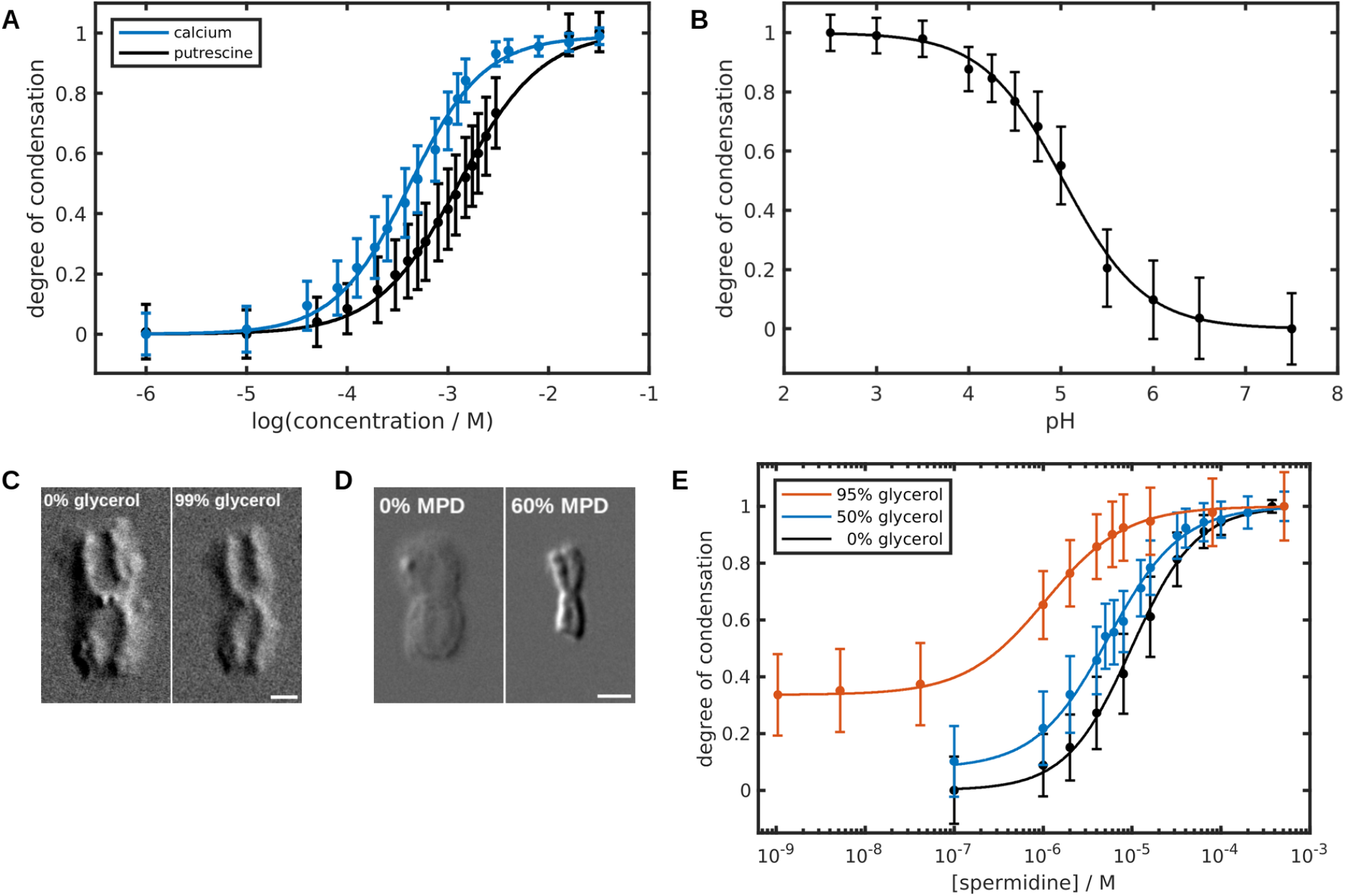
Validation of predictions from polyelectrolyte gel theory: pH and solvent effects. (A) Condensation profiles for chromosomes titrated with putrescine (black; data from Figure 3A) or calcium chloride (blue), indicating the existence of ion-specific effects. (B) The degree of condensation is governed by the fractional dissociation of ionizable groups intrinsic to chromatin. Chromosomes were condensed by stepwise reduction of the pH from 7.5 to 2.5. The normalized degree of condensation, averaged for *n* = 205 chromosomes and fit to the Hill model, is plotted as a function of the solution pH. (C) Chromosomes exposed to 0% (left) or 99% (right) glycerol, and imaged by DIC (indicated percentages are volume/volume). The negligible increase in contrast reflects both partial condensation and the elevated background refractive index (*n* ≈ 1.47 for 99% glycerol compared to *n* ≈ 1.33 for water). (D) Chromosomes exposed to 0% (left) or 60% (right) 2-methyl-2,4-pentanediol (MPD), and imaged by DIC (indicated percentages are volume/volume). Note that the dielectric constants for pure glycerol and MPD at 298 K are *ε* = 42.5 and *ε* = 25.1 (*97*), respectively, compared to *ε* = 78.3 for water. The degree of condensation (and evidently the Flory-Huggins parameter for the chromatin-solvent system) increases inversely with the dielectric constant of the solvent. Scale bars for panels C and D represent 2 µm. (E) Condensation profiles obtained by titrating spermidine in the presence of different concentrations of glycerol show that glycerol potentiates the effect of spermidine. Curves represent averages of *n* = 367 (0% glycerol), *n* = 188 (50% glycerol), and *n* = 223 (95% glycerol) chromosomes. The black curve (0% glycerol) is also displayed in Figure 3A.

**Figure S6:**
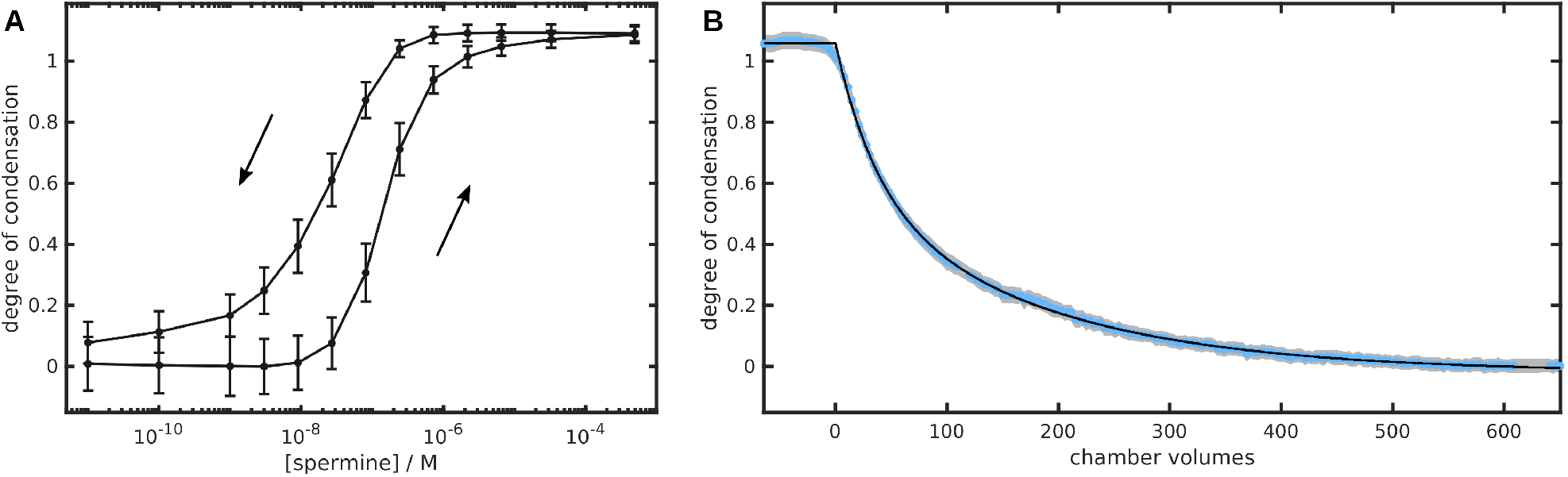
Chromosome condensation is hysteretic. (A) Hysteresis of chromosome condensation, observed by varying the concentration of spermine. Chromosomes were condensed by spermine and subsequently decondensed by its withdrawal (path indicated by the arrows), resulting in a hysteresis loop (*n* = 617). The degree of condensation (normalized with respect to the degrees of condensation produced by solutions containing 0 and 375 µM spermidine) is plotted as a function of the spermine concentration. During each step of decondensation, chromosomes were imaged after flowing at least 500 chamber volumes of solution. (B) Chromosomes condensed by treatment with 4 mM spermine were decondensed by flowing a solution consisting of 5 mM Tris (pH 7.5) and 2 mM KCl. The degree of condensation of *n* = 150 chromosomes (blue points; gray shading denotes the 99% confidence interval) was measured as a function of the volume of buffer administered, normalized by the volume of the flow chamber. The black curve represents a biexponential fit of the data.

**Figure S7:**
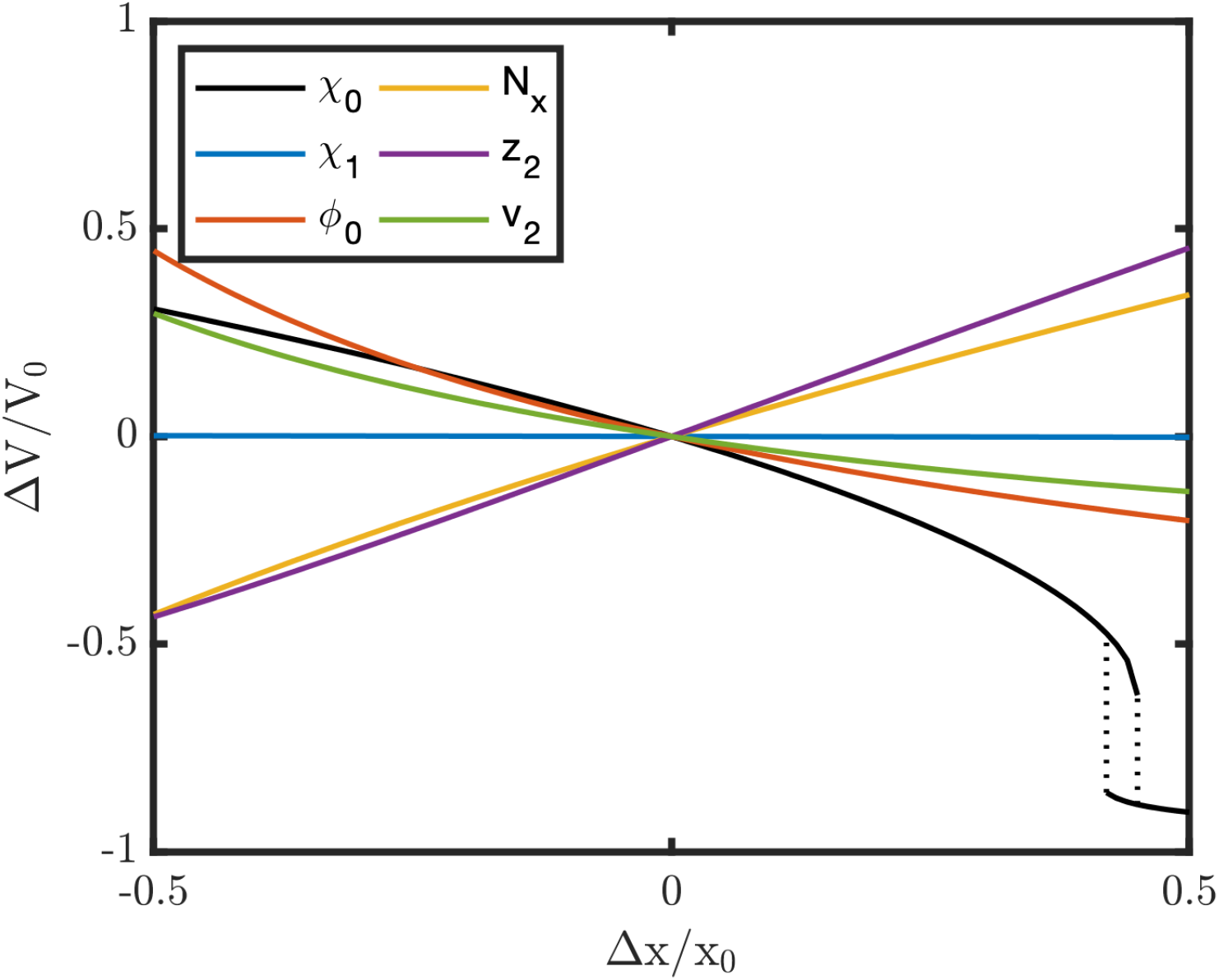
Effect of varying model parameters on gel volume. Effect of variation of individual model parameters (x) on gel volume (V) at physiologic salt (125 mM monovalent salt). The ordinate represents the difference in gel volume relative to that of the starting volume (ΔV*/*V_0_), and the abscissa represents the relative difference of parameters (Δx*/*x_0_), which are shown in the legend (*χ*_0_, *χ*_1_, *φ*_0_, *N_x_*, *z*_2_, and v_2_). The initial values of the parameters and volume are those for a gel parameterized as in Figure 3C (solid lines). A VPT is predicted to occur upon raising *χ*_0_, indicated by the dotted lines that form a hysteresis loop. Our model does not consider coupling between *χ* and fixed charge density (*z*_2_).

**Movie S1: Liberation of the scaffold by DNase digestion of a decondensed chromosome.** Differential interference contrast time-lapse recording of a decondensed chromosome exposed to deoxyribonuclease I.

